# Heterogeneous subpopulations of GABAAR-responding neurons coexist in physiological and pathological mature neuronal networks at increasing scales of complexity

**DOI:** 10.1101/2023.05.25.542321

**Authors:** I. Colombi, M. Rastogi, M. Parrini, M. Alberti, M. Chiappalone, M. Nanni, A. Contestabile, L. Cancedda

## Abstract

GABA is the main inhibitory neurotransmitter in adults. Depolarizing/excitatory GABA responses have been well characterized at the level of neuronal-population *average* during typical *neurodevelopment* and partially in pathology. However, no investigation has specifically assessed whether a *mosaicism* of cells with either depolarizing/excitatory or hyperpolarizing/inhibitory GABAergic responses exists in *adult* animals in health/disease. Here, we showed that such mosaicism is present both in adult WT and Down syndrome (DS) mice, as assessed at increasing scales of neuronal-network complexity (cultures, brain-slices, behaving mice). Nevertheless, WT mice presented a lower percentage of cells with depolarizing GABA than DS mice. Restoring the mosaicism of hyperpolarizing and depolarizing GABA-responding neurons to WT levels rescued anxiety behaviour in DS mice. We also found heterogeneous GABAergic responses in mature control and trisomic human iPSC-derived neurons. Thus, a heterogeneous population of GABA-responding cells exists in physiological/pathological conditions in mature mouse and human neurons, possibly contributing to disease-associated behaviours.

## INTRODUCTION

γ-Aminobutyric acid (GABA) is the main inhibitory neurotransmitter in the adult mammalian brain by acting on chloride (Cl^-^)-permeable GABAA receptors (GABAARs). In mature neurons, the activity of the Cl^-^ exporter KCC2 predominates over that of the Cl^-^ importer NKCC1, resulting in a low intracellular chloride concentration ([Cl^-^]i). Therefore, the chloride reversal potential through GABAARs (EGABA) lies below the cellular resting membrane potential (RMP) in mature neurons. This makes inward the flow of negatively charged chloride ions through the receptor; thus, GABAAR signalling is hyperpolarizing and inhibitory ^1–3^ in adult animals. Conversely, during neuronal development (until the first or second postnatal week in rodents), GABA exerts a depolarizing and possibly excitatory effect due to low expression of KCC2 in immature neurons, which leads to a high [Cl^-^]i and an outward flow of Cl^-^ through GABAARs^2, 4^. Depolarizing GABAergic signalling during development is fundamental for organizing early patterns of neuronal activity by enabling neurons to fire and thus form their first synaptic connections ^5–11^. The excitatory-to-inhibitory developmental switch in the polarity of GABAAR signalling has been very well characterized at the level of the neuronal population average ^12–14^. However, there is still a possibility that a small subpopulation of neurons with depolarizing polarity of GABAAR-mediated responses may persist even after full neuronal maturation. While this small percentage of depolarizing GABA neurons could be hidden within the population average or may be fairly interpreted as a biological outlier or as standard noise in experimental datasets present in the literature, these neurons may in fact represent a specific subpopulation of cells coexisting with the large majority of cells with hyperpolarizing GABAAR-mediated signalling and have important physiological and pathological roles.

The rapidly increasing evidence reported in the last twenty years of literature has indicated an altered NKCC1/KCC2 expression ratio and depolarizing GABAAR-mediated signalling as a common feature of a very large number of brain pathologies ^14–16^. These brain pathologies range from neurodevelopmental disorders (e.g., Down syndrome, Rett syndrome, Fragile X syndrome, 22q11.2 microdeletion syndrome, and schizophrenia) to neurodegenerative disorders (e.g., Huntington disease and Parkinson’s disease) and neurological conditions (e.g., chronic pain and some forms of epilepsy ^14–16^). However, whether the NKCC1/KCC2 expression ratio in these brain disorders reflects generally depolarized GABAAR-mediated responses in most neurons or simply a specific increase in the numerical extent of a subpopulation of neurons with depolarizing responses is not known, although it may have important functional implications.

Here, by performing electrophysiological and/or Ca^2+^ imaging recordings on large populations of neurons at increasing scales of network complexity (*in vitro* neuronal cultures, *ex vivo* brain slices and *in vivo* freely behaving mice), we found a significant population of mature neurons with NKCC1-dependent, depolarizing GABAergic responses both in adult wild-type (WT) mice and in a widely studied mouse model of DS (Ts65Dn adult mice). Treatment with the NKCC1 inhibitor bumetanide fully rescued the numerical extent of the population of neurons with inhibitory GABAergic signalling and normalized both anxiety-related behaviours and paradoxical responses to the pharmacological activation of GABAARs with a clinically used anxiolytic drug in adult DS animals without altering the behaviour of adult WT animals. Finally, we found a heterogeneous population of mature neurons with mixed responses to GABAAR signalling also in human isogenic control and DS neurons derived from induced pluripotent stem cells (iPSCs) obtained from a person with a form of mosaic trisomy 21. Additionally, in human mature neurons, trisomic networks showed a significantly larger subpopulation of neurons with depolarizing GABAergic signalling than that in the corresponding isogenic control neuronal networks. Thus, a heterogeneous population of GABA-responding mature neurons is present in human and murine neuronal networks under physiological and pathological conditions, where it possibly plays a role in dysfunctional behaviors.

## RESULTS

### A mixed population of neurons with hyperpolarizing or depolarizing GABA signalling is present in mature WT and Ts65Dn cultures *in vitro*

To investigate whether a heterogeneous population of mature neurons with mixed (hyperpolarizing and depolarizing) GABA-mediated responses is present *in vitro*, we performed Ca^2+^ imaging experiments in *bona fide* mature (21 days *in vitro*, DIV) hippocampal neuronal primary cultures from WT (B6EiC3) and Ts65Dn mice. Ts65Dn mice are a well-studied animal model of DS in which the average depolarizing GABAergic signalling has already been described by electrophysiological recordings in both mature neuronal cultures and acute brain slices from adult animals ^17, 18^. To benchmark our results and to evaluate the natural decay of depolarizing GABA responses during development ^19–21^, we also imaged neurons at earlier stages of maturation (2, 7, and 15 DIV). In particular, we evaluated Ca^2+^ responses elicited by bath application of GABA (100 μM) in WT and Ts65Dn neuronal cultures loaded with the Ca^2+^-sensitive dye Fluo-4 (Fig. 1A). Indeed, the transient increase in intracellular neuronal calcium concentration ([Ca^2+^]i) due to the opening of voltage-gated Ca^2+^ channels by the application of GABAAR agonists can be used as a proxy to assess the depolarizing action of GABA, which has been previously described using this approach for neuronal population *averages* during development ^11, 21, 22^. By calculating the percentage of cells showing GABA-induced Ca^2+^ responses in our neuronal cultures at increasing DIV, we found that not all neurons underwent the expected switch in the GABA response from excitatory to inhibitory, even at the latest stage analysed (Fig. 1B). Indeed, at 21 DIV (well beyond the completion of the GABA developmental switch ^20, 21, 23^), we still observed a population of WT neurons (14% of cells) presenting depolarizing GABA signalling. This population was significantly smaller than that in immature WT neurons at 2 DIV (80% of neurons with depolarizing GABA) or at 7 DIV (26%) but similar to that found at 15 DIV (15%; Fig. 1B). Interestingly, while the initial developmental drop (2-7 DIV) in the population of neurons with depolarizing GABA-mediated responses in Ts65Dn neurons was similar to that of WT neurons, the population of Ts65Dn neurons with depolarizing GABA responses was significantly greater at later time points (15 and 21 DIV; Fig. 1B). These results were corroborated by the significantly higher levels of NKCC1 expression (and no difference in KCC2 levels) that indeed we found in Ts65Dn neurons at 15 and 21 DIV in comparison to WT controls (Supplementary Fig. 1). Next, to specifically assess the involvement of NKCC1 in the population of neurons retaining depolarizing responses to GABA in WT and Ts65Dn cultures, we evaluated the percentage of cells showing GABA-induced Ca^2+^ responses in the presence of the NKCC1 inhibitor bumetanide (10 μM ^17^) in neuronal cultures at 15 DIV (Fig. 1C and D). Fifteen DIV is a stage when GABAergic signalling has already switched from excitatory to inhibitory in most WT neurons, and it is when we first detected a significant difference in the developmental trajectory of GABA-mediated responses in comparison to Ts65Dn neurons. We found that bumetanide treatment strongly reduced the percentage of neurons showing GABA-induced Ca^2+^ responses in both WT and Ts65Dn cultures (Fig. 1D). Accordingly, when we performed live imaging experiments in primary hippocampal neurons at 15 DIV with the chloride-sensitive dye MQAE, we found that WT neurons displayed a bimodal distribution in the raw MQAE fluorescence values, indicating the presence of neurons with both low and high values of [Cl^-^]i, even at this advanced stage of development (Fig. 1E and F). Conversely, the distribution of MQAE values of Ts65Dn neurons was broader and shifted towards lower values (reflecting a higher [Cl^-^]i) compared with that of WT cultures, as indicated by both the beanplot and the boxplot quantifications (Fig. 1F and 1G). Interestingly, NKCC1 inhibition by bumetanide (10 μM) abolished the bimodal distribution of the raw MQAE values and significantly decreased [Cl^-^]i (*i.e.,* higher MQAE fluorescence) in WT neurons (Fig. 1E, F and G). Similarly, NKCC1 inhibition decreased the higher [Cl^-^]i found in Ts65Dn neurons to within more physiological levels, similar to those of WT neurons (Fig. 1E, F and G).

**Figure 1.**
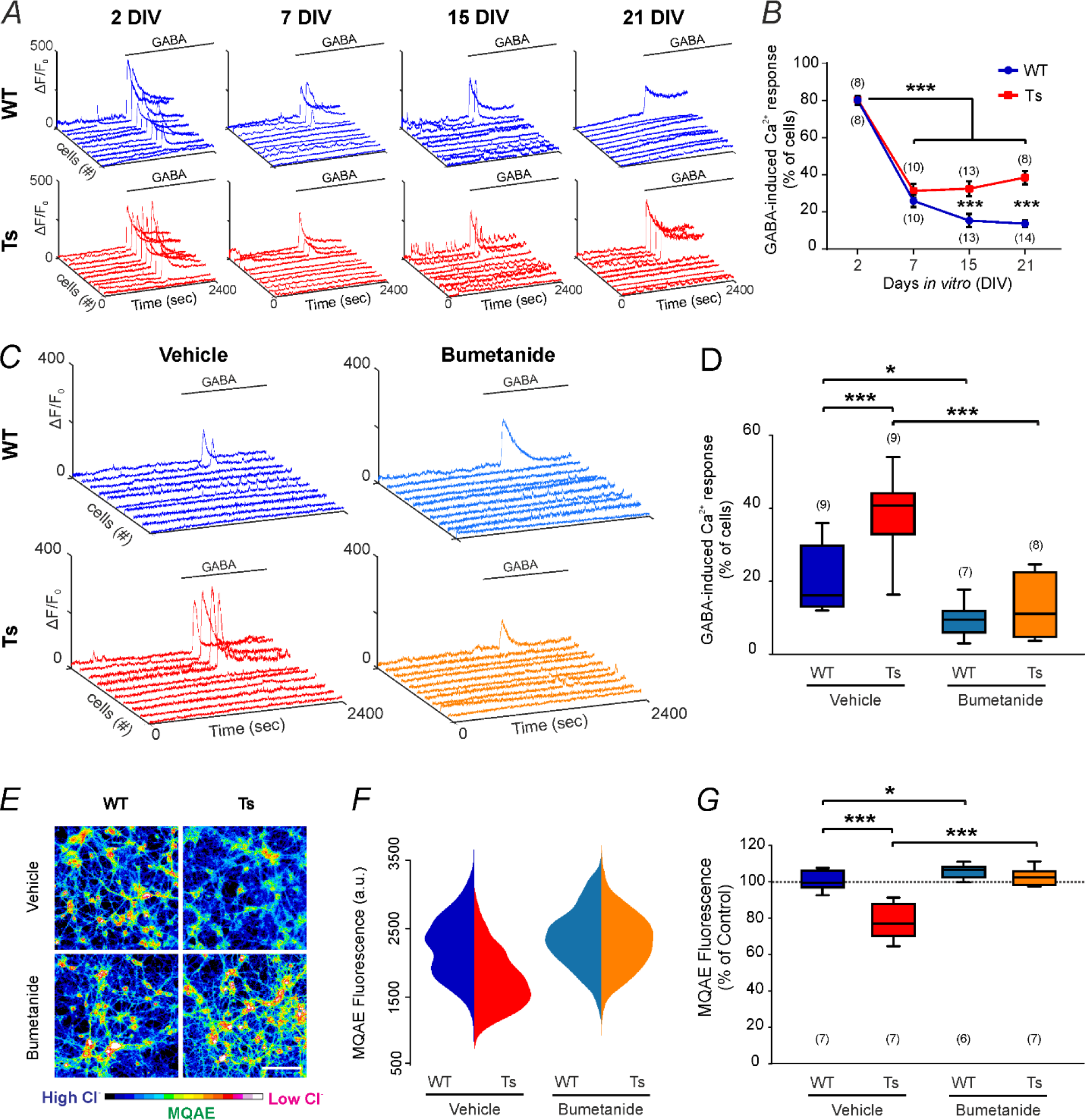
Mixed subpopulations of neurons with depolarizing or hyperpolarizing GABA signalling are present during the *in vitro* development of neuronal primary cultures. A) Representative calcium traces obtained from live imaging of cultured WT and Ts65Dn hippocampal neurons loaded with the Ca^2+^- sensitive dye Fluo-4 upon bath application of GABA (100 μM) at different time points during *in vitro* development. **B)** Quantification of the mean percentage (± SEM) of neurons showing GABA-induced Ca^2+^ responses (*i.e.*, depolarizing GABA responses) in experiments as in A. Numbers in parentheses indicate the number of analysed coverslips for each time point (obtained from 3-4 independent neuronal cultures). ***P<0.001, Tukey post hoc test following two-way ANOVA. **C)** Representative calcium traces of neurons pretreated with vehicle (0.01% DMSO) or the NKCC1 inhibitor bumetanide (10 μM) upon bath application of GABA at 15 DIV. **D)** Quantification of the percentage of neurons showing depolarizing GABA responses in experiments as in C. Boxplots indicate median and 25th-75th percentiles, whiskers represent the 5th- 95th percentiles. The numbers in parentheses indicate the number of analysed coverslips for each experimental group (obtained from 3-5 independent neuronal cultures). *P<0.05, ***P<0.001, Tukey *post hoc* test following two-way ANOVA. **E)** Representative images of WT and Ts65Dn hippocampal neurons during imaging experiments with the chloride-sensitive dye MQAE upon treatment with vehicle (0.01% DMSO) or bumetanide (10 μM) at 15 DIV. The fluorescence intensity of the dye (colour-coded at the bottom) is inversely proportional to [Cl^-^]i. Scale bar: 100 μm. **F)** Beanplot showing the distribution of MQAE raw fluorescence values for all neurons imaged as in E (WT-vehicle 840 neurons, Ts-vehicle 818 neurons, WT-bumetanide 597 neurons, Ts-bumetanide 840 neurons). **G)** Quantification of the average [Cl^-^]i with MQAE in the same dataset described in F. Boxplots indicate the median and 25th-75th percentiles, and whiskers represent the 5th-95th percentiles. The numbers in parentheses indicate the number of analysed coverslips for each experimental group (obtained from 3 independent neuronal cultures). *P<0.05, ***P<0.001, Tukey *post hoc* test following two-way ANOVA.

Finally, to confirm that the presence of a subpopulation of neurons with depolarizing GABA signalling in mature neurons was indeed mediated through GABAARs, we performed control experiments in WT and Ts65Dn neuronal cultures at 15 DIV in the presence of the specific GABAA agonist muscimol (10 μM), the GABAAR antagonist bicuculline (100 μM) or the GABABR antagonist CGP55845 (10 μM ^17^). We found that muscimol elicited GABAAR-dependent depolarizing Ca^2+^ responses completely similar to those elicited by GABA application (Supplementary Fig. 2A and B). This effect was inhibited by bumetanide application both in WT and Ts65Dn neurons (Supplementary Fig. 2A and B). Accordingly, GABA-induced Ca^2+^ responses were largely abrogated by blocking GABAARs with bath application of bicuculline (Supplementary Fig. 2C and D). Of note, pretreatment of neurons with the L-type voltage-gated calcium channel (CaV1) blocker nifedipine (10 μM) also strongly reduced GABA-induced Ca^2+^ responses, indicating that GABA-induced depolarization generates a Ca^2+^ inflow mainly through the activation of CaV1 channels in both WT and Ts65Dn neurons (Supplementary Fig. 2 C and D). Finally, GABA-induced Ca^2+^ responses were enhanced after blocking GABABRs with CGP55845 bath application, consistent with the inhibitory function of GABABRs on neuronal networks.

Together, these data indicated the presence of a heterogeneous population of neurons with hyperpolarizing or NKCC1-mediated depolarizing GABAA signalling in mature WT cultures, with a larger subpopulation of GABA-depolarizing neurons in mature Ts65Dn cultures.

### Subpopulations of neurons with functionally heterogeneous GABAAR-mediated responses are present in mature WT or Ts65Dn neuronal networks cultured over MEAs

To assess the functional consequences of the presence of the two disproportional subpopulations of neurons with hyperpolarizing or depolarizing GABAergic responses in WT and Ts65Dn mature neuronal networks, we recorded their firing activity by microelectrode arrays (MEAs). This technique allows the simultaneous recording of the activity from diverse spots of a large population of neurons without interfering with neuronal [Cl^-^]i. In the first set of experiments, we assessed the neuronal mean firing rate (MFR) after blocking endogenous GABAAergic signalling by bath application of bicuculline (20 μM ^17^) on WT and Ts65Dn (21 DIV) cultures, which were previously preincubated with vehicle (0.01% DMSO) or bumetanide (10 μM, Fig. 2A). For each electrode, we evaluated the changes in activity elicited by bicuculline by calculating the MFR ratio (MFR after the treatment over the MFR during baseline). As expected, the average MFR ratio of WT control cultures (pretreated with vehicle; Fig. 2B, dark blue box; raster plots in Supplementary Fig. 3A and B) in the presence of bicuculline was (5 times) higher than the MFR of untreated neurons (dotted line). Similarly, WT cultures pretreated with bumetanide (light blue box) also displayed a large average MFR ratio, which was even greater than that seen in the corresponding vehicle-treated WT cultures. Conversely, the Ts65Dn control cultures (pretreated with vehicle; Fig. 2B, red box; raster plots in Supplementary Fig. 3 B) showed a significantly lower average MFR ratio, indicating a greatly blunted response to GABAAR inhibition by bicuculline, when compared to that in WT cultures. Interestingly, pretreatment with bumetanide restored the full response to bicuculline application in Ts65Dn neurons, similar to levels observed in WT cultures (Fig. 2B, orange box; raster plots in Supplementary Fig. 3 B).

**Figure 2.**
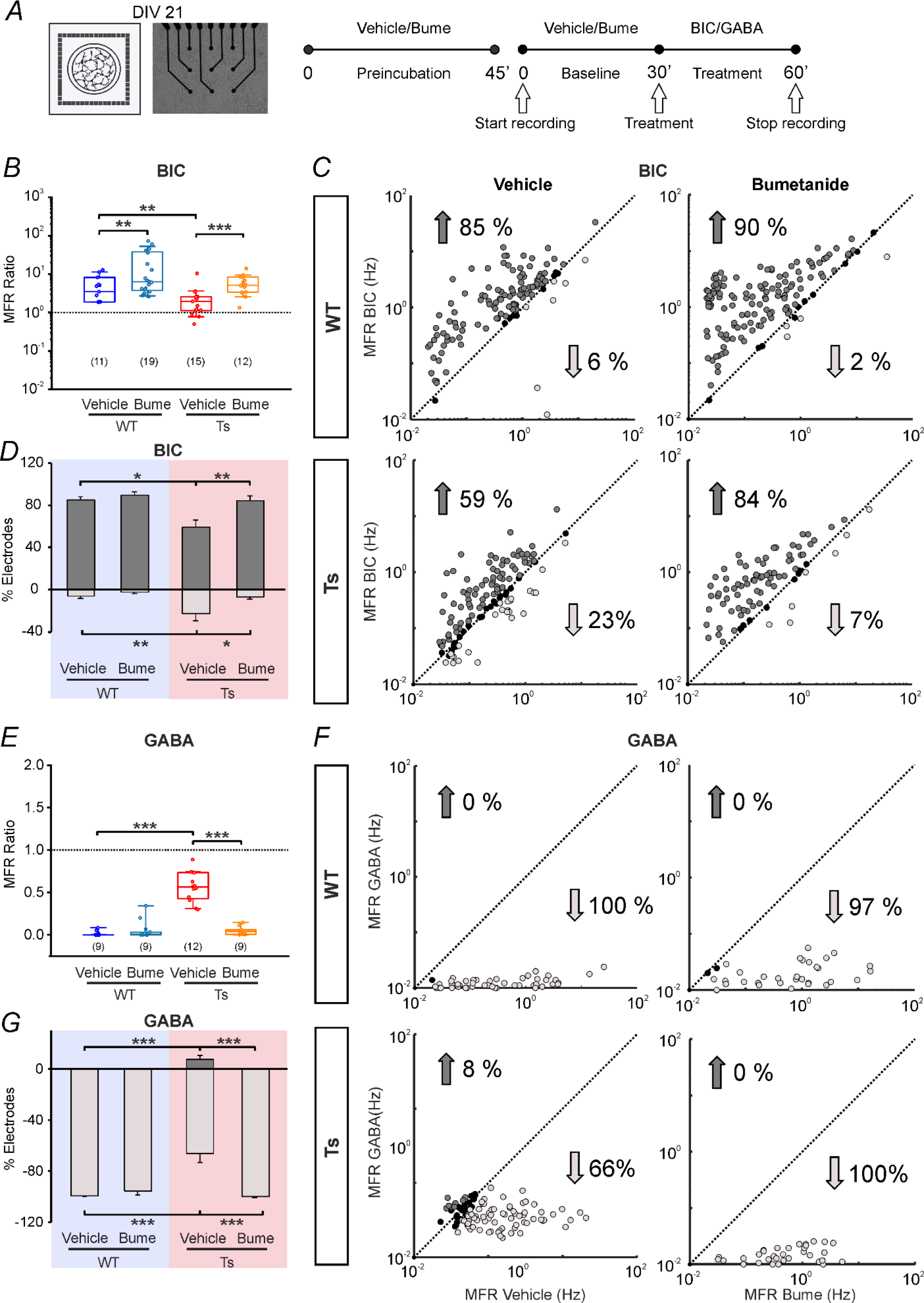
Mixed subpopulations of neurons with hyperpolarizing or depolarizing GABAAR signalling are present in mature neuronal cultures grown over MEAs. A) *Left*: Schematic representation of a primary hippocampal neuronal culture grown over a microelectrode array (MEA) for electrophysiological recordings. *Centre*: Representative transmitted-light image showing an MEA seeded with hippocampal neurons at 21 DIV. *Right*: Schematic representation of the experimental protocol; neuronal cultures were preincubated with vehicle (0.01% DMSO) or bumetanide (10 μM) for 45 minutes followed by 30-minute recording of spontaneous activity. Neurons were then recorded for an additional 30 minutes 10 minutes after the addition of bicuculline (BIC, 20 μM) or GABA (100 μM). **B, E)** Quantification of the mean firing rate (MFR) ratio of the WT and Ts65Dn neuronal cultures upon BIC (B) or GABA (E) bath application. The MFR ratio over the baseline firing (dotted line) was calculated for each electrode and then averaged for each MEA. An MFR ratio higher than 1 (representing the baseline level) indicates an increase in activity upon treatment, whereas a ratio below 1 indicates a decrease in activity. In the boxplot, the small square indicates the mean, the central line illustrates the median, the box limits indicate the 25th and 75th percentiles, the whiskers represent the 5th-95th percentiles, and each dot represents the MFR ratio for each recorded culture. The numbers in parentheses indicate the number of analysed MEAs for each experimental group (obtained from 5 independent neuronal cultures). **p<0.01; *** p<0.001; Tukey’s *post hoc* test following two-way ANOVA. **C, F**). Scatter plots showing the MFR for each active electrode (plotted as a dot) from all recorded MEAs (in B or E) seeded with WT and Ts65Dn neurons before (x-axis) and after (y-axis) bath application of BIC (C) or GABA (F). Dark grey dots represent electrodes showing a significant increase in the MFR. Light grey dots represent electrodes showing a significant decrease in the MFR. Black dots represent electrodes showing no significant changes in the MFR. Significant changes in the MFR (numbers with arrows) for each electrode upon BIC or GABA application were evaluated by bootstrap analysis. **D, G)** Quantification of the average percentage number (± SEM) of MEA electrodes (in the same experiments in B, C, E and F), showing significant changes in the MFR by bootstrap analysis after BIC (D) or GABA (G) administration in comparison to their basal conditions in WT (blue) and Ts65Dn (pink) neuronal cultures. *P<0.05, **P<0.01, *** p<0.001; Tukey’s *post hoc* test following two-way ANOVA on ranked transformed data.

As we hypothesized the presence of a heterogeneous population of neurons with hyperpolarizing or depolarizing GABAAR-mediated responses in WT and Ts65Dn cultures, we took advantage of the large number of MEA observation points (*i.e.,* electrodes) scattered through the network to extrapolate information about possibly different subpopulations of neurons ^24, 25^. For each active (*i.e.,* mean firing rate higher than 0.02 spikes/sec) electrode, we evaluated whether the MFR upon bicuculline application was significantly different from the basal level of activity by using a bootstrap method. Both the scatterplot representation of single-electrode MFR activity before and after bicuculline application (Fig. 2C) and the average percentage of electrodes that were significantly changed by bootstrap analysis (Fig. 2D) clearly showed an increase in the firing activity for the majority of the electrodes (85% of MFR-increasing electrodes) for vehicle-treated WT cultures. Notably, in the same recordings, we found that 6% of electrodes demonstrated a significant decrease in the firing rate activity, consistent with the presence of a subpopulation of neurons presenting depolarizing rather than hyperpolarizing GABA signalling in mature WT cultures. Bumetanide treatment in these WT cultures caused a small (and nonsignificant) change in the percentage of electrodes showing either an increase or decrease in MFR activity (Fig. 2C and D, Supplementary Fig. 3B). This likely reflects only a very subtle enhancement of inhibitory GABAergic drive upon bumetanide application due to the prevalence of KCC2 expression over NKCC1 expression at this stage of development ^26, 27^.

Conversely, vehicle-treated Ts65Dn mature cultures showed far more heterogeneous responses to the blockade of GABAAR signalling by bicuculline than WT cultures demonstrated. Indeed, we found a higher percentage of electrodes that decreased MFR activity (23% of MFR-decreasing electrodes) and a decrease in electrodes showing increased MFR (59% of MFR-increasing electrodes) in comparison to the WT mature cultures (Fig. 2C), consistent with alterations in GABAAR-mediated responses towards more depolarizing values (Fig. 2D). Notably, pretreatment of Ts65Dn cultures with bumetanide significantly shifted the distribution of the MFR values to WT levels (84% of MFR-increasing electrodes and 7% of MFR-decreasing electrodes, Fig. 2C), indicating a full rescue of GABAAR-mediated inhibition (Fig. 2D). We found no statistically significant difference in the percentage of electrodes that did not respond to the treatment of both WT and Ts65Dn mature cultures under all conditions (Supplementary Fig. 4 A and B).

Next, we analysed the average MFR variation upon treatment with exogenously applied GABA (100 μM ^17^; Fig. 2E). Mature WT cultures pretreated with either vehicle or bumetanide at 21 DIV showed a strong decrease in the average MFR ratio (well below 1, Fig. 2E, raster plots in Supplementary Fig. 3C), consistent with the expected decrease in the neuronal activity level upon GABA application. Conversely, vehicle-treated Ts65Dn cultures displayed a significantly higher average MFR ratio and an overall much larger variability among MEAs than controls upon GABA treatment. Interestingly, pretreatment with bumetanide fully restored inhibitory GABAergic signalling in trisomic cultures to WT levels (Fig. 2E, raster plots in Supplementary Fig. 3C).

As described above, we used bootstrap methods to evaluate the difference in the firing rate activity at the single-electrode level. Scatterplot representation of single MFR values for each active electrode and the average percentage of electrodes that significantly changed by bootstrap analysis before and after GABA application in vehicle-treated WT cultures showed an abrupt suppression of the firing activity in virtually all (99.7% of MFR-decreasing electrodes) active electrodes (Fig. 2F and G, Supplementary Fig. 3C). Bumetanide pretreatment had no significant effect on WT cultures (Fig. 2F and G; Supplementary Fig. 3C). In vehicle-treated Ts65Dn cultures with exogenously applied GABA, the distribution of the single MFR values was clearly different from that observed for WT cultures, with a wide dispersion of the data. Only 68% of electrodes were characterized by a decreased MFR, and 8% of electrodes showed an increased MFR, indicating a general reduction in the efficacy of GABA inhibition and even partial GABA-driven excitatory activity (Fig. 2F and G, Supplementary 3C). Notably, treatment with bumetanide fully restored inhibitory GABA signalling in Ts65Dn cultures, as demonstrated by the distribution and average of single MFR values, which was indistinguishable from that of WT cultures (100% of MFR-decreasing electrodes, Fig. 2F and G, Supplementary 3C). In support to the above data indicating reduced inhibition in Ts65Dn neurons, we found a significant difference in the number of electrodes that did not change significantly upon GABA application between vehicle-treated Ts65Dn and WT mature cultures (24% *vs.* 0% of electrodes with no MFR changes, respectively; Supplementary Fig. 4C). This effect was also completely rescued by bumetanide treatment (Supplementary Fig. 4C).

To strengthen our findings, we also performed experiments with the GABAAR positive allosteric modulator (PAM) diazepam (a benzodiazepine widely used as an anxiolytic medication) in WT and Ts65Dn mature neuronal cultures (Supplementary Fig. 5A). We found that diazepam application (1 μM) consistently modulated network activity by decreasing firing in most (but not all) recorded electrodes in WT cultures pretreated with either vehicle (4% of MFR-increasing electrodes, 77% of MFR-decreasing electrodes) or bumetanide (3% of MFR-increasing electrodes, 78% of MFR-decreasing electrodes; Supplementary Fig. 5B, C, D and E). Conversely, in Ts65Dn cultures, the MFR distribution was more dispersed, with 11% of the electrodes showing increased activity and 63% of the electrodes showing decreased activity. Notably, NKCC1 inhibition by bumetanide fully recovered diazepam-mediated GABAergic inhibition, as indicated by the average MFR ratio, MFR distribution (5% of MFR-increasing electrodes and 78% of MFR-decreasing electrodes) and average percentage of electrodes that were significantly changed by bootstrap analysis of Ts65Dn mature neurons, which were similar to those of WT mature cultures (Supplementary Fig. 5B, C, D and E).

Overall, these results demonstrated the presence of a heterogeneous population of neurons in both WT and Ts65Dn mature cultures characterized by differences in the polarity of GABAAR-mediated responses. In particular, Ts65Dn neuronal cultures showed a larger subpopulation of cells characterized by GABAAR- mediated depolarization, which depends on NKCC1 activity.

### Subpopulations of neurons with functionally heterogeneous GABAAR-mediated responses are present in hippocampal slices from adult WT and Ts65Dn mice recorded by MEAs

Next, we assessed whether a mixed population of neurons with heterogeneous GABA-mediated responses was also present *ex vivo*. To this aim, we prepared acute hippocampal brain slices (which preserve at least in part the original architecture of the brain circuits) from adult (10-12-week-old) WT and Ts65Dn mice and recorded them by MEAs (Fig. 3A).

**Figure 3.**
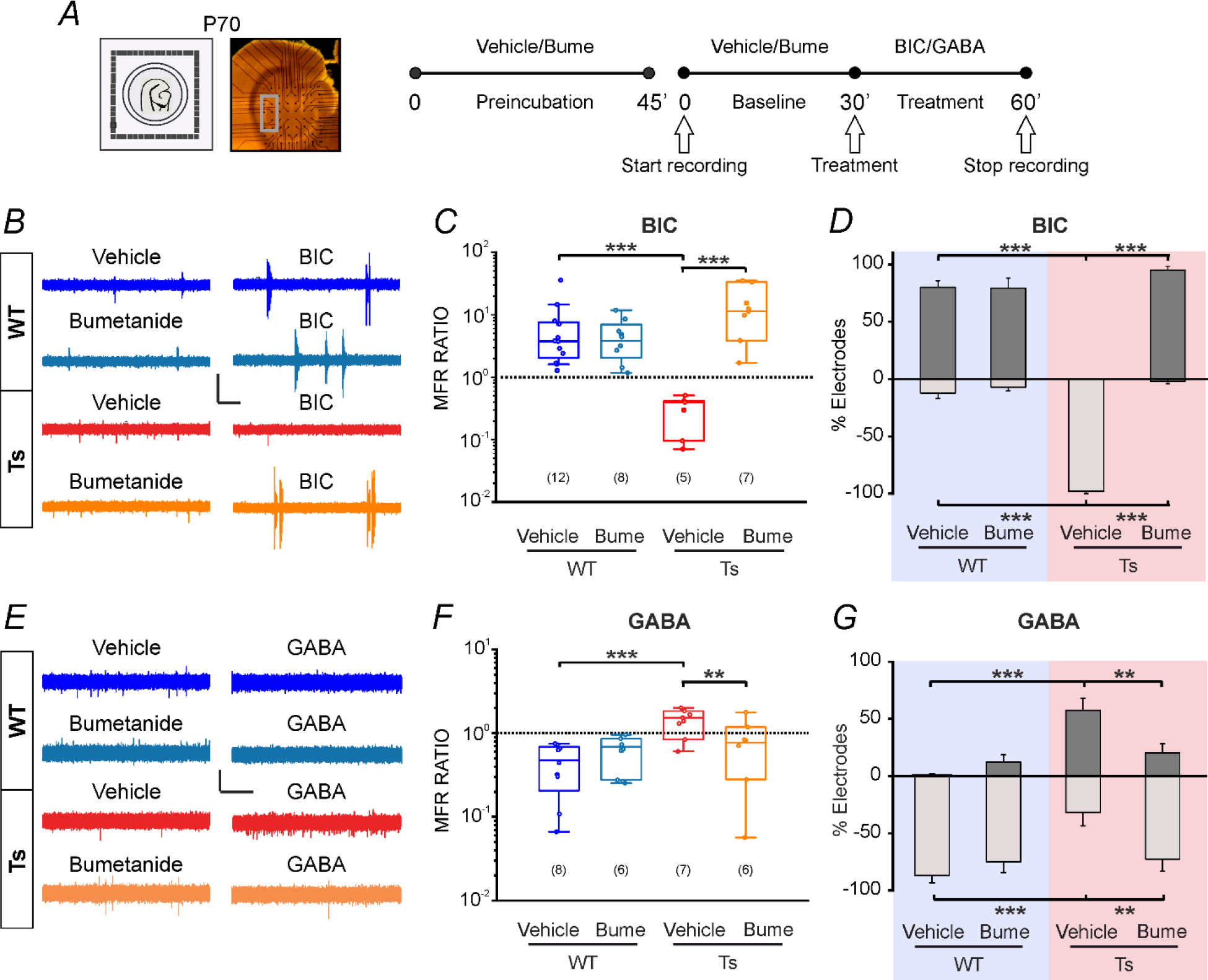
Mixed subpopulations of neurons with hyperpolarizing or depolarizing GABA signalling are present in acute hippocampal slices from adult WT and Ts65Dn mice. A) *Left*: Schematic representation of an acute brain slice of the hippocampus-entorhinal cortex (EC) region from an adult animal (postnatal day (P)70). The slice is positioned on an MEA for electrophysiological recordings; *Centre*: Representative picture of the experimental setup with MEA electrodes and an overlying hippocampus-EC slice. The grey square highlights the analysed electrodes, which were positioned in the CA1 region. *Right*: Schematic representation of the experimental protocol. Brain slices were first preincubated with vehicle (0.01% DMSO) or bumetanide (10 μM) for 45 minutes. Then, we recorded 30 minutes of spontaneous activity in the presence of vehicle or bumetanide followed by 30 minutes of recording 10 minutes after the addition of bicuculline (BIC; 20 μM) or GABA (100 μM). **B, E)** Representative high-pass filtered electrophysiological traces (>300 Hz) for WT and Ts65Dn slices pretreated with vehicle or bumetanide and recorded before and after bath application of BIC (B) or GABA (E). Scale bars, (B): 80 μV, 2 s; (E): 30 μV, 2 s. **C, F)** Quantification of the neuronal mean firing rate (MFR) ratio from WT and Ts65Dn slices upon BIC (C) or GABA (F) bath application. The MFR ratio over the baseline firing (dotted line) was calculated for each electrode and then averaged for each MEA. An MFR ratio higher than 1 (representing the baseline level) indicates an increase in activity upon the treatment, whereas a ratio below 1 indicates a decrease in activity. In the boxplot, the small square indicates the mean, the central line illustrates the median, the box limits indicate the 25th and 75th percentiles, the whiskers represent the 5th-95th percentiles, and each dot represents the MFR ratio for each recorded slice. The numbers in parentheses indicate the number of analysed slices for each experimental group. **p<0.01; *** p<0.001; Tukey’s *post hoc* test following two-way ANOVA. **D, G)** Quantification of the average percentage number ± SEM of MEA electrodes (in the same experiments in C and F), showing changes of at least 15% in the MFR after BIC (D) or GABA (G) administration in WT (blue) and Ts65Dn (pink) slices. **p<0.01; *** p<0.001; Tukey’s *post hoc* test following two-way ANOVA on ranked transformed data.

In the first set of experiments, we inhibited endogenous GABAergic signalling by bath application of bicuculline (20 μM^17^). In particular, after preincubation with vehicle (0.01% DMSO) or bumetanide (10 μM ^17^) for 45 minutes, we recorded 30 minutes of baseline data (at the level of the CA1 hippocampal region) followed by an additional 30 minutes of recording after bicuculline application (Fig. 3A). As expected, the average MFR ratio of both vehicle-treated and bumetanide-treated WT slices clearly showed a large increase in the firing rate activity upon bicuculline application (Fig. 3B and C). Conversely, bicuculline administration to vehicle-treated Ts65Dn slices resulted in a clear decrease in the average MFR ratio in comparison to the basal conditions (Fig. 3B and C). Remarkably, Ts65Dn slices pretreated with bumetanide showed a pronounced increase in the MFR ratio and reached the levels observed in WT slices (Fig. 3B and C). This outcome is in line with a possibly large subpopulation of NKCC1-dependent GABA depolarizing neurons in Ts65Dn adult mice and with data obtained by cell-attached patch-clamp recordings in the literature ^17, 18^.

Next, to specifically assess whether a heterogeneous population of neurons with mixed responses to *endogenous* GABA is also present in adult WT and Ts65Dn brain slices, we quantified the percentage of electrodes showing significant changes in spiking activity upon bicuculline application in comparison to their baseline level. We found that the majority of the electrodes detected a significantly increased firing rate upon blocking GABAARs with bicuculline in WT slices (80% of MFR-increasing electrodes), irrespective of vehicle or bumetanide pretreatment (Fig. 3D). Remarkably, we found a significant percentage of electrodes in WT slices that decreased the firing rate upon bicuculline treatment, reflecting the presence of a subpopulation of neurons with depolarizing GABA signalling (12% of MFR-decreasing electrodes). Bumetanide pretreatment caused a small positive shift in the number of electrodes with decreasing spiking activity, but the difference did not reach statistical significance (Fig. 3D). Strikingly, in vehicle-treated Ts65Dn slices, almost all the active electrodes showed a decrease in the average MFR level upon bicuculline administration (0% of MFR-increasing electrodes, 97% of MFR-decreasing electrodes), indicating a very profound alteration of inhibitory GABAergic signalling. Notably, Ts65Dn slices pretreated with bumetanide displayed a high MFR at all the active electrodes, with the percentage of electrodes showing an increase in the MFR ratio comparable to that of WT slices (95% of MFR-increasing electrodes, 2% of MFR-decreasing electrodes), indicating a complete rescue of inhibitory GABAergic signalling (Fig. 3D). We found no differences in the percentage of electrodes that did not detect a response to bicuculline among all experimental groups (Supplementary Fig. 6 A and B).

Next, we also analysed the firing activity variation upon application of *exogenous* GABA (100 μM ^17^) to brain slices of WT and Ts65Dn adult mice. As expected, vehicle-treated WT slices clearly showed a decrease of the average MFR (Fig. 3E and F). Bumetanide pretreatment only led to a nonsignificant shift towards higher MFR ratio values (Fig. 3E and F). Conversely, vehicle-treated Ts65Dn control slices showed an increased average MFR ratio upon GABA administration in comparison to that in WT slices (Fig. 3E and F). Notably, pretreatment with bumetanide fully restored inhibitory GABAergic signalling in Ts65Dn slices, as indicated by the clear decrease in the average MFR ratio to levels similar to those of WT slices (Fig. 3E and F).

Again, when we evaluated changes in the MFR variation upon GABA application at the single-electrode level, we found mixed subpopulations of GABA-responding neurons in both WT and Ts65Dn slices, with the difference between WT and Ts65Dn slices even more profound than that in neuronal cultures. Indeed, exogenously applied GABA caused a decrease in the firing rate activity at most (but not all) of the electrodes in vehicle-treated WT slices (1% of increasing of MFR-increasing electrodes; 87% of MFR- decreasing (Fig. 3G). Notably, vehicle-treated Ts65Dn slices displayed a pronounced mixed response to GABA treatment with 32% of electrodes that showed a decrease in the MFR and 57% of those that showed increased MFR activity (Fig. 3G). Notably, Ts65Dn slices pretreated with bumetanide showed an evident negative shift in the percentage of electrodes that detected significantly changed responses. Indeed, the MFR at most of the electrodes decreased upon GABA application (20% of MFR-increasing electrodes and 72% of MFR-decreasing electrodes) similar to the WT slices and indicative of a large recovery of physiological proportions between the neuronal subpopulations with hyperpolarizing or depolarizing GABAergic signalling. We found no difference in the percentage of electrodes that did not respond to GABA in any of the experimental groups (Supplementary Fig. 6C).

Of note, we also evaluated a second model of DS (the Dp(16)1Yey/+ mice ^25^) and found no difference in the subpopulations of mature neurons with heterogeneous GABAAR-mediated responses in comparison to their corresponding WT littermates (C57BL/6J). Indeed, slices from adult Dp(16)1Yey/+ mice showed a mixed population of neurons responding to GABA with increased or decreased MFRs but to levels very similar to those of their corresponding WT littermates (Supplementary Fig. 7A, B, C and D). Therefore, the increased population of neurons with depolarizing GABAA-mediated responses that we identified in adult Ts65Dn animals was not replicated in a second mouse model of DS. However, hippocampal NKCC1 levels in adult Dp(16)1Yey/+ animals were similar to those of WT animals, further strengthening the NKCC1 dependency of depolarizing GABAergic responses in Ts65Dn mice (Supplementary Fig. 7E, F and G).

Overall, these results demonstrated the presence of a heterogeneous population of neurons characterized by differences in GABAAR and NKCC1-mediated responses in *ex vivo* slices from adult WT and Ts65Dn mice that appeared even larger than those in mature neuronal cultures.

### Subpopulations of neurons with heterogeneous GABAAR-mediated responses recorded by *in vivo* Ca^2+^ imaging are present in adult freely moving WT and Ts65Dn mice

Next, to further strengthen our results, we performed Ca^2+^ imaging experiments in a network of neurons *in vivo*. For these experiments, we chose Ca^2+^ imaging because it allows the activity of a large population of neurons to be recorded with a high degree of spatial resolution in freely moving mice ^28^. Moreover, we chose to modulate GABAergic activity *in vivo* with the positive GABAAR allosteric modulator diazepam (instead of GABA) because it is an FDA-approved drug with translational validity, has recognized behavioural outcomes and is known to readily pass the blood‒brain barrier. We also discarded the usage of GABAAR blockers (which we used in fact in culture and slice experiments), as we reasoned that they could potentially induce epileptic activity *in vivo.* This would have strongly complicated the analysis of Ca^2+^ signals and the interpretation of the results especially in the case of DS animals, which have higher susceptibility to seizures ^17, 29^.

To perform Ca^2+^ imaging experiments in the hippocampus *in vivo*, we implanted a microendoscopic probe coupled to a miniaturized head-mounted microscope in freely moving, adult (3-4 months old) WT and Ts65Dn mice previously injected in the hippocampus with an adeno-associated virus (AAV) to express the Ca^2+^ indicator GCaMP6f in dorsal CA1 pyramidal neurons under the control of the neuronal CamK2a promoter ^30^ (Fig. 4A). We set up a crossover study in which the same animals were first imaged after administration of vehicle (2% DMSO in saline, i.p.) followed by diazepam (2 mg/kg, i.p.); after two weeks, imaging was performed again upon administration of bumetanide (0.2 mg/kg, i.p.) followed by diazepam (Fig. 4B). This experimental design allowed us to longitudinally follow the same neurons across the two experimental sessions, evaluating the changes in activity induced by the two pharmacological manipulations (diazepam and bumetanide) in each cell, and thus comparing the effect of diazepam with and without bumetanide pretreatment at the single-cell level. Finally, to avoid potential photobleaching and/or phototoxicity during imaging, we adopted an intermittent imaging protocol ^31^ (Fig. 4B).

**Figure 4.**
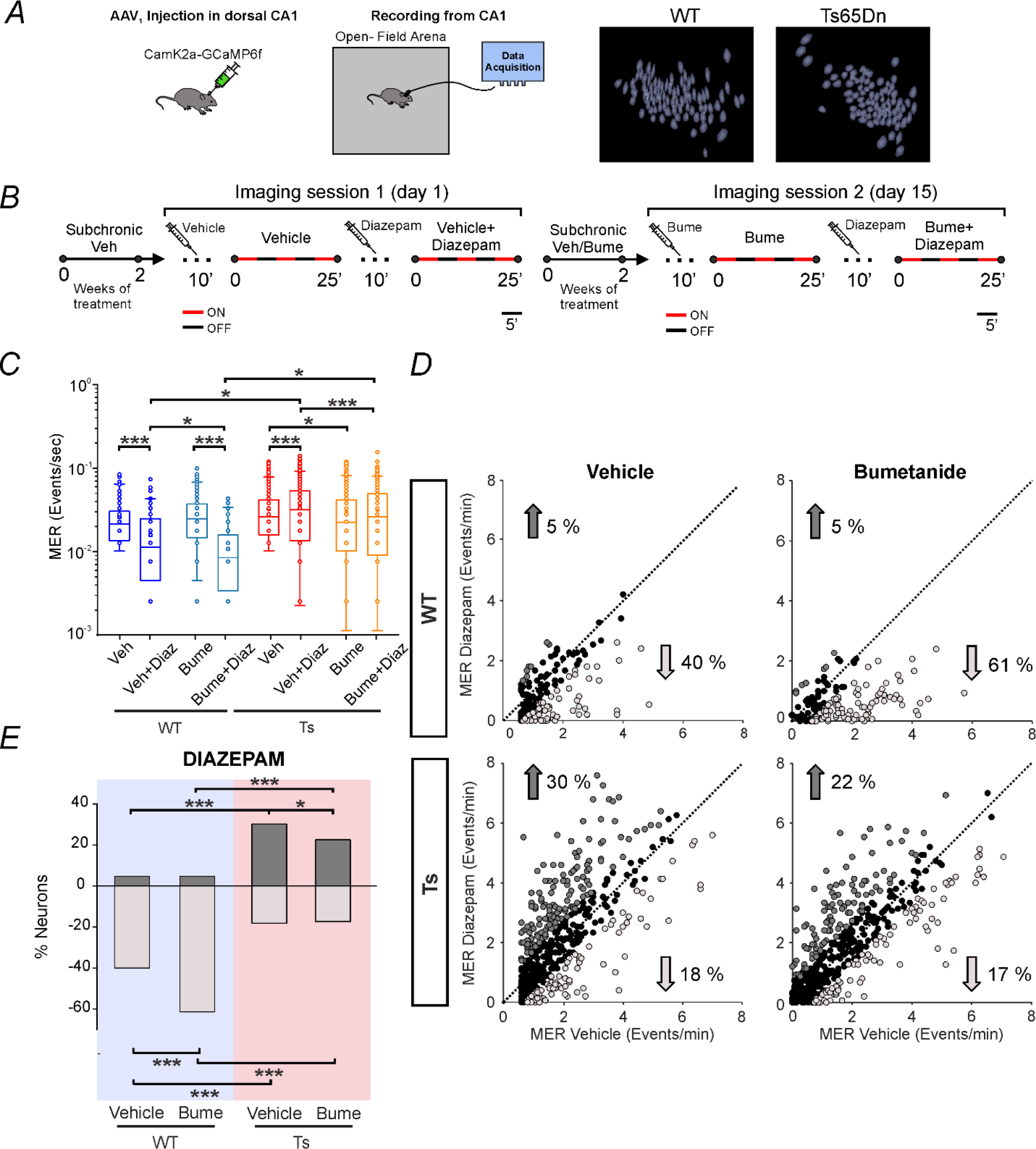
Bumetanide treatment differentially affects neuronal activity *in vivo* in the hippocampus of freely moving adult WT and Ts65Dn mice and partially rescues aberrant GABAAR signalling in Ts65Dn animals. A) Schematic representation of the experimental setup. *Left:* Two- to three-month-old WT and Ts65Dn mice received a stereotaxic injection in the dorsal hippocampal CA1 region with AAV viruses expressing the Ca^2+^-sensor GCaMP6f under the control of the CamK2a promoter. Ts65Dn and WT mice were then implanted with a microendoscopic probe, and neuronal activity was longitudinally assessed by recording *in vivo* Ca^2+^ events within the CA1 hippocampal region of freely moving mice with a miniaturized head-mounted microscope. *Right:* Examples of projection maps of recorded neurons in a WT and a Ts65Dn animal. **B)** Schematic representation of the experimental protocol timeline. Ca^2+^ events were imaged in the same neuron in two consecutive sessions before and after administration of the GABAAR positive allosteric modulator diazepam (2 mg/kg), following a subchronic (2 weeks) treatment with either vehicle (2% DMSO in saline) or bumetanide (0.2 mg/kg). Within each imaging session, neuronal activity was recorded in periods of 5 minutes (“ON”, red) alternated to 5 minutes of rest (“OFF”, blue) to collect 15 min of neuronal data for each session while avoiding phototoxicity. **C)** Quantification of single neuron mean event rates (MERs) of WT and Ts65Dn mice before and after diazepam administration in experiments as in B. In the boxplot, the small square indicates the mean, the central line illustrates the median, the box limits indicate the 25th and 75th percentiles, and whiskers represent the 5th-95th percentiles. Each dot indicates the binned value of the MER (bin size=0.005 events/sec) obtained from individual neurons (from 5 WT and 8 Ts65Dn adult mice). *p<0.05; *** p<0.001; Tukey’s *post hoc* test following a linear mixed effects model (with two factors). **D)** Scatter plots showing the MER for each active (MER>0.01 events/s in vehicle) neuron (plotted as a dot) in C before (x-axis) and after (y-axis) administration of diazepam. Dark grey dots represent neurons showing a significant increase in the MER. Light grey dots represent neurons showing a significant decrease in the MER. Black dots represent neurons showing no significant changes in the MER. Significant changes in the MER (number with arrows) for each neuron upon diazepam application were evaluated by bootstrap analysis. **E)** Quantification of the percentage of neurons (in the same experiments in C), showing significant changes in the MER by bootstrap analysis after diazepam administration in comparison to their basal conditions in WT (blue) and Ts65Dn (pink) mice. * p<0.05, *** p<0.001, Chi-Square test with Sidak adjustment for multiple comparisons.

As expected, the mean Ca^2+^ event rate (MER) in adult WT mice significantly decreased upon diazepam administration after pretreatment with vehicle. Notably, bumetanide pretreatment led to a further significant decrease in MER upon diazepam treatment in adult WT mice, suggesting the presence of a subpopulation of NKCC1-dependent GABA-depolarizing neurons also *in vivo* in freely moving animals. In agreement with the cell culture and brain slice data, bumetanide application *per se* did not cause any significant variation in the MER of WT animals (Fig. 4C). Interestingly, Ts65Dn neurons showed a significant mean MER increase upon diazepam treatment. This indicates that alteration of GABAAR-dependent inhibition and possibly the depolarizing action of GABA also occurs *in vivo* in adult DS mice, as already reported *in vitro* ^17, 32, 33^. Remarkably, bumetanide pretreatment reduced the baseline MER and prevented the diazepam-induced MER increase in Ts65Dn animals (Fig. 4C), indicating an NKCC1 dependency in the aberrant response to diazepam in adult DS mice.

Next, we specifically investigated the presence of subpopulations of neurons with heterogeneous GABA responses also in freely moving adult animals by bootstrap methods to compute the percentage of neurons that significantly changed their activity by diazepam treatment. The majority of responding neurons of vehicle-treated WT animals showed significantly lowered frequencies of Ca^2+^ transients following diazepam administration (40% MFR-decreasing neurons), indicating the mainly inhibitory action of GABA (Fig. 4D and E; raster plots in Supplementary Fig. 8A and B). Notably, a small neuronal subset (5%) in vehicle-treated WT animals had a significantly increased MER, reflecting the presence of a subpopulation of pyramidal CA1 neurons showing depolarizing responses to GABAAR signalling also *in vivo* in WT animals. However, a remarkable 55% of WT neurons did not show a significant change (Supplementary Fig. 8C) in response to diazepam *in vivo*, indicating a general low inhibitory chloride driving force in neurons *in vivo*. Anyhow, bumetanide pretreatment in WT animals caused a significant enhancement of inhibitory GABAAR signalling *in vivo*, as shown by the higher percentage of neurons that had a decreased MER (61% MFR-decreasing neurons; Fig. 4D and E; Supplementary Fig. 8A and B) and the significant decrease in nonresponding neurons (Supplementary Fig. 8C).

Vehicle-treated Ts65Dn animals displayed a much wider dispersion of single MER values *in vivo* than WT animals (Fig. 4D and E). In particular, we found that only 18% of neurons showed a decreased MER and 30% of neurons showed an increased MER upon diazepam treatment, indicating a strong reduction in the efficacy of GABAAR-dependent inhibition and even paradoxical GABAAR-driven depolarizing activity (Figure 4 D and E, Supplementary Fig. 8B). Notably, Ts65Dn mice pretreated with bumetanide showed a significantly lower percentage of neurons that increased the MER upon treatment with diazepam (22% of MER-increasing neurons, Fig. 4D and E, Supplementary Fig. 8B) and an increased number of nonresponding neurons (60% of MER-nonchanging neurons, Supplementary Fig. 8C), indicating that blocking NKCC1 activity rescues GABAAR-dependent inhibition in a subpopulation of Ts65Dn pyramidal neurons *in vivo*.

Overall, these results demonstrated the presence of a heterogeneous population of hyperpolarizing or depolarizing GABAAR-responding neurons in adult WT and Ts65Dn animals even *in vivo*, with proportions comparable to *ex vivo* data, but with much higher percentages of GABAAR nonresponding neurons.

### Heterogeneous subpopulations of neurons with distinct responses to bumetanide or diazepam are present in WT and Ts65Dn mice *in vivo*

Since bumetanide pretreatment reduced the average baseline neuronal Ca^2+^ MER and prevented, on average, the diazepam-induced neuronal MER increase in adult Ts65Dn animals, we next investigated these subpopulations of bumetanide-responsive cells in more detail. Conveniently, our longitudinal recordings allowed us to follow the Ca^2+^ activity of the same neuron throughout the bumetanide and diazepam treatments. Therefore, we first investigated whether the neurons that showed a decrease in MER upon bumetanide administration were also resistant to diazepam-induced stimulation, demonstrating that increased NKCC1 activity mediates the paradoxical effects of diazepam in mature Ts65Dn neurons. To this aim, we calculated the variation in Ca^2+^ activity induced by bumetanide over vehicle treatment (Ratio 1, Fig. 5A) and the variation caused by diazepam *plus* bumetanide treatment over diazepam alone (Ratio 2, Fig. 5A) for each Ts65Dn neuron. To evaluate possible relationships between changes in neuronal activity induced by the two treatments, we plotted the value of Ratio 1 *vs.* Ratio 2 (*i.e.,* the variation in neuronal activity induced by bumetanide *vs.* diazepam treatments) for each neuron and divided the obtained scatterplot into 4 different quadrants, reflecting the different possible neuronal categories determined by changes in activity induced by the two drugs (Fig. 5B). Specifically, cells showing a decrease in MER after both bumetanide and bumetanide *plus* diazepam treatments were classified into category 1; cells showing an increase in MER after bumetanide and a decrease after bumetanide *plus* diazepam administrations were classified into category 2; cells showing an increase in MER after both bumetanide and bumetanide *plus* diazepam administrations were classified into category 3; and cells showing a decrease in MER after bumetanide and an increase after bumetanide *plus* diazepam administration were classified into category 4 (Fig. 5B). We found a significant positive correlation (P<0.001, Spearman’s test; Fig. 5C) for the activity of Ts65Dn neurons upon bumetanide and diazepam application. Notably, the majority of cells were allocated to category 1, thus implying the presence of a bumetanide-responsive population of Ts65Dn neurons in which NKCC1 inhibition can ameliorate the paradoxical effects induced by diazepam treatment (Fig. 5D).

**Figure 5.**
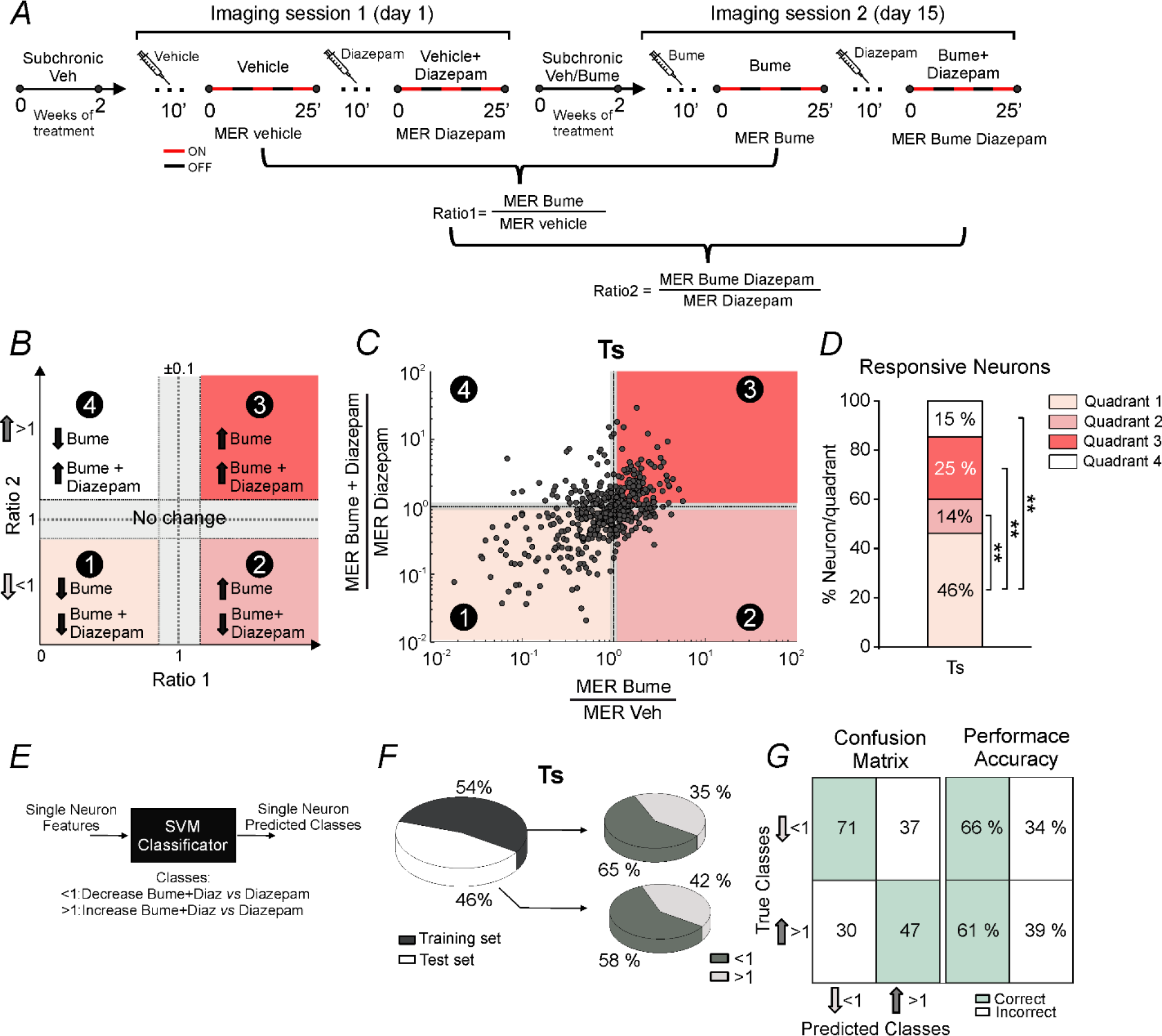
Mixed subpopulations of neurons with hyperpolarizing or depolarizing GABAAR signalling respond differently to bumetanide administration *in vivo*. A) *S*chematic representation of the experimental protocol timeline (top) together with representation of the analysis approach (bottom). **B)** Schematic representation of the possible distribution of neuronal subpopulations based on MER variation in response to bumetanide or diazepam treatment (Ratio 1 *vs.* Ratio 2, see main text). Cells showing MER ratio changes below 10% were excluded from the analysis (shadowed grey area). The numbers in the black dots indicate the four quadrants representing the possible different responses (increasing or decreasing arrows) of neuronal activity upon diazepam or bumetanide treatment over that elicited after vehicle treatment. **C)** Scatterplot showing the neuronal population distribution based on MER variation in response to bumetanide or diazepam treatment for Ts65Dn neurons in the same experiments described in Figure 4. Pearson correlation showed a significant positive correlation between the responses to bumetanide or diazepam treatment in neurons from Ts65Dn mice. **D)** Quantification of the percentage of neurons for each quadrant on the same dataset described in C. ** p<0.01 Chi-square test with Sidak adjustment for multiple comparisons. **E)** Schematic representation of the support vector machine (SVM) classifier used for discrimination between the two classes of variation (*i.e.,* decrease (<1) or increase (>1) of activity after bumetanide *plus* diazepam when compared to vehicle *plus* diazepam). **F)** *Left*: Percentage of Ts65Dn neurons used as a training set or a test set for the SVM classification. *Right*: The upper pie chart reports the percentage of neurons that fell into the two categories (>1 and <1) for the training set. The lower pie chart reports the percentage of neurons that fell in the two categories (>1 and <1) after running the SVM classification on the test set. **G)** *Left*: Confusion matrix showing the performance accuracy (correct green and incorrect white) of the SVM classifier in Ts65Dn neurons based on the two populations of diazepam-responding neurons (<1 decrease upon diazepam *plus* bumetanide or >1 increase upon bumetanide *plus* diazepam when compared to diazepam *plus* vehicle). *Right*: Percentage of neurons in the test set correctly (green boxes) or incorrectly (white boxes) classified into the two categories (<1 or >1).

To further strengthen this finding, we used machine-learning algorithms (support vector machine classifier, SMV; Fig. 5E) to try to predict the diazepam response elicited in each Ts65Dn cell based on the knowledge of the previous response caused by bumetanide administration to the same cell (*i.e.*, to understand whether the response to bumetanide could predict the future response to diazepam in terms of variation of Ca^2+^ MER). For simplicity, we used only two different categories that referred to either an increase (>1) or a decrease (<1) in the MER during diazepam *plus* bumetanide treatment when compared to diazepam alone (*i.e.,* changes in Ratio 2; Fig. 5E). First, we split the Ca^2+^ MER *in vivo* dataset into one training and one test set, with percentages of neurons equal to 54% and 46%, respectively, and with a balanced representation of the categories (*i.e.,* Ratio 2>1 and Ratio 2<1; Fig. 5F). Next, we used the training set to feed the SVM for the classification of neurons in the test set. When we evaluated the performance accuracy of the predictive model using a confusion matrix ^34^ on the test set, we found that 71 out of the 108 total neurons belonging to the <1 category and 47 out of 77 total neurons belonging to the >1 category were correctly classified into the two categories (Fig. 5G, left). Conversely, 37 neurons were wrongly discriminated as the >1 category instead of the <1 category, and 30 neurons were assigned to the <1 category instead of the >1 category (Fig. 5G, left). By computing the percentage of neurons per category, we successfully discriminated 66% of neurons in the <1 category and 61% of neurons in the >1 category (Fig. 5G, right; Supplementary Table 1).

Taken together, these findings suggested that different levels of NKCC1 activity—probed by diverse responses to bumetanide—are possibly responsible for the paradoxical effect of diazepam on neuronal activity in mature Ts65Dn cells *in vivo*.

### Diazepam treatment exerts abnormal behavioural effects in adult Ts65Dn mice, which are rescued by bumetanide treatment

Next, we investigated the functional implications of our findings at the behavioural level. To this aim, we first treated adult (2-3 months old) WT and Ts65Dn mice with vehicle (2% DMSO in saline, i.p.) or bumetanide (0.2 mg/kg, i.p.) ^17, 35^ followed by saline (0.9% NaCl, i.p.) or diazepam (2 mg/kg, i.p.) administration 30 minutes later and assessed anxiety behaviours and the anxiolytic effects of diazepam in three different behavioural tests after an additional 30 minutes (Fig. 6A). In the dark-light test, vehicle *plus* saline-treated Ts65Dn mice showed a reduction in the time spent in the illuminated (*i.e.,* the potentially unsafe) area of the apparatus compared to WT mice, indicative of increased anxiety. Notably, this behaviour was fully rescued by bumetanide treatment, indicating a possible role for NKCC1 in the altered behaviour of adult Ts65Dn mice in this test. We also found a trend towards increased time in the light zone in WT animals upon bumetanide treatment, but this outcome did not reach statistical significance. Furthermore, in agreement with the larger population of cells responding with increased activity to diazepam that we found in imaging experiments in adult Ts65Dn mice, treatment with diazepam actually worsened the anxiety behaviour of Ts65Dn mice rather than rescuing it. Interestingly, pretreatment with bumetanide completely rescued anxiety behaviour in Ts65Dn animals even after diazepam administration (Fig. 6B). We found no differences in the total number of transitions between the dark and light zones for WT and Ts65Dn mice (Fig. 6C), indicating that general activity was not altered by drug treatments.

**Figure 6.**
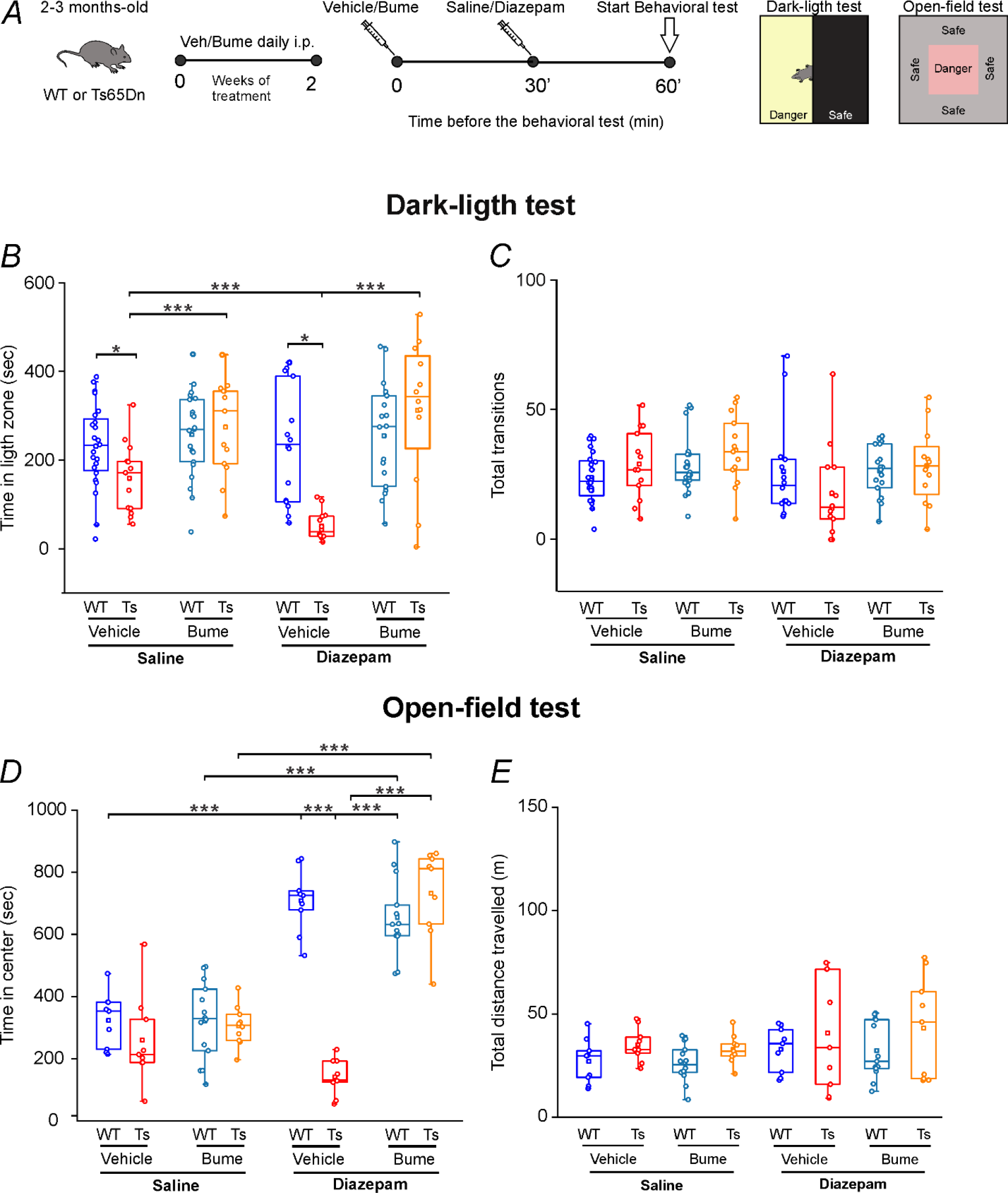
Bumetanide treatment rescues aberrant behavioural responses to the benzodiazepine diazepam in adult Ts65Dn mice. A) Schematic representation of the protocol for the diverse experimental groups. Adult WT and Ts65Dn mice were treated with bumetanide (0.2 mg/kg i.p.) or the corresponding vehicle (2% DMSO in saline) for two weeks and then assessed in two different anxiety tests. On the day of testing, mice were pretreated with bumetanide or vehicle and then tested 30 minutes after diazepam (2 mg/kg i.p.) or saline administration. **B)** Quantification of time spent in the light zone of the dark-light test for WT and Ts65Dn mice. **C)** Quantification of the total number of transitions between the two zones of the dark-light test for WT and Ts65Dn mice. **D)** Quantification of the time spent in the centre of the arena during the open-field test for WT and Ts65Dn mice. **E)** Quantification of the total distance travelled in the open field test for WT and Ts65Dn mice. In all boxplots, the small square indicates the mean, the central line illustrates the median, the box limits indicate the 25th and 75th percentiles, whiskers represent the 5th-95th percentiles, and each dot indicates a value obtained from individual animals. *** p<0.001; Tukey’s *post hoc* test following two-way ANOVA or two-way ANOVA on ranked transformed data.

Next, we evaluated the effect of diazepam treatment in the open-field test, another behavioural task widely used to assess anxiety behaviour in mice. We found that diazepam administration in adult WT animals strongly increased the time spent in the central portion (*i.e.,* the potentially unsafe area) of the open-field arena, indicative of the well-known anxiolytic effect of the drug. Vehicle *plus* saline-treated adult Ts65Dn mice showed a trend towards increased anxiety compared to WT mice in this test (slight decrease in time spent in the centre of the arena). Additionally, diazepam administration showed a trend towards increased anxiety in Ts65Dn mice (Fig. 6D). Nevertheless, none of these trends reached statistical significance (Fig. 6D). Pretreatment with bumetanide had a very large effect and fully rescued the anxiolytic effect of diazepam in adult Ts65Dn mice to WT levels, consistent with the results of the dark-light test. We found no differences in the total distance travelled for either WT or Ts65Dn mice (Fig. 6E), again indicating no potentially confounding effects of the drugs on locomotor activity in this test.

In line with the recent literature ^36^, we also found a substantial lack of effect of diazepam administration in adult Ts65Dn mice in the elevated plus maze test, as indicated by the similar percentage of time spent in the closed arms after saline or diazepam administration (Supplementary Fig. 9 A, B). Conversely, diazepam-treated WT mice showed a decrease in the percentage of time spent in the closed arms, reflecting the expected anxiolytic effect of the drug. Notably, in agreement with the results from the open-field test, pretreatment of adult Ts65Dn mice with bumetanide significantly decreased the percentage of time spent in the closed arms and simultaneously increased the percentage of time spent in the open arms (Supplementary Fig. 9 A, B). Thus, bumetanide pretreatment was able to restore the anxiolytic effect of diazepam in Ts65Dn animals. Although diazepam administration caused a significant reduction in the average total number of arm entries for both WT and Ts65Dn mice (Supplementary Fig. 9C), this outcome was not paralleled by a decrease in the total distance travelled (Supplementary Fig. 9D), indicating that the other effects observed upon diazepam treatment did not likely depend on decreased locomotor activity.

Together, these data revealed that diazepam treatment induced atypical responses, ranging from a paradoxical anxiogenic effect to a lack of efficacy in Ts65Dn mice compared to that in WT adult mice in diverse behavioural tasks. Most importantly, bumetanide treatment fully re-established the anxiolytic effect of diazepam in all behavioural tasks in DS animals.

### A mixed population of neurons with depolarizing or hyperpolarizing GABA signalling is present in human iPSC-derived neurons from control and trisomic cell lines from a person with mosaic DS

To further strengthen and increase the translational value of our findings, we investigated whether a heterogeneous population of mature neurons with mixed responses to GABAAR signalling was also present in human cells. To this aim, we evaluated neurons derived from iPSCs from a previously established DS line (DS4) and its corresponding isogenic control cell line (DS2U). Both cell lines derived from a same person with a mosaic form of DS ^37^. In particular, we evaluated Ca^2+^ responses elicited by bath application of GABA (100 μM) in mature (60 days after plating from the final differentiation ^20, 38^) human DS and control neurons. By calculating the percentage of cells showing GABA-induced Ca^2+^ responses, we found that a subpopulation of neurons (14% of cells) presented depolarizing GABA responses also in human control mature cell cultures (Fig. 7A and B). This population was significantly larger in DS neuronal cultures at the same developmental stage (62% of GABA-responding cells). Interestingly, treatment with bumetanide (10 μM) significantly reduced the percentage of neurons showing GABA-induced Ca^2+^ responses in DS cultures without significantly altering the control neurons (Fig. 7A and B). These results were corroborated by the significantly higher levels of NKCC1 expression (and no significant difference in KCC2 expression) that we found in mature DS neurons in comparison to control neurons (Fig. 7 C, D and E).

**Figure 7.**
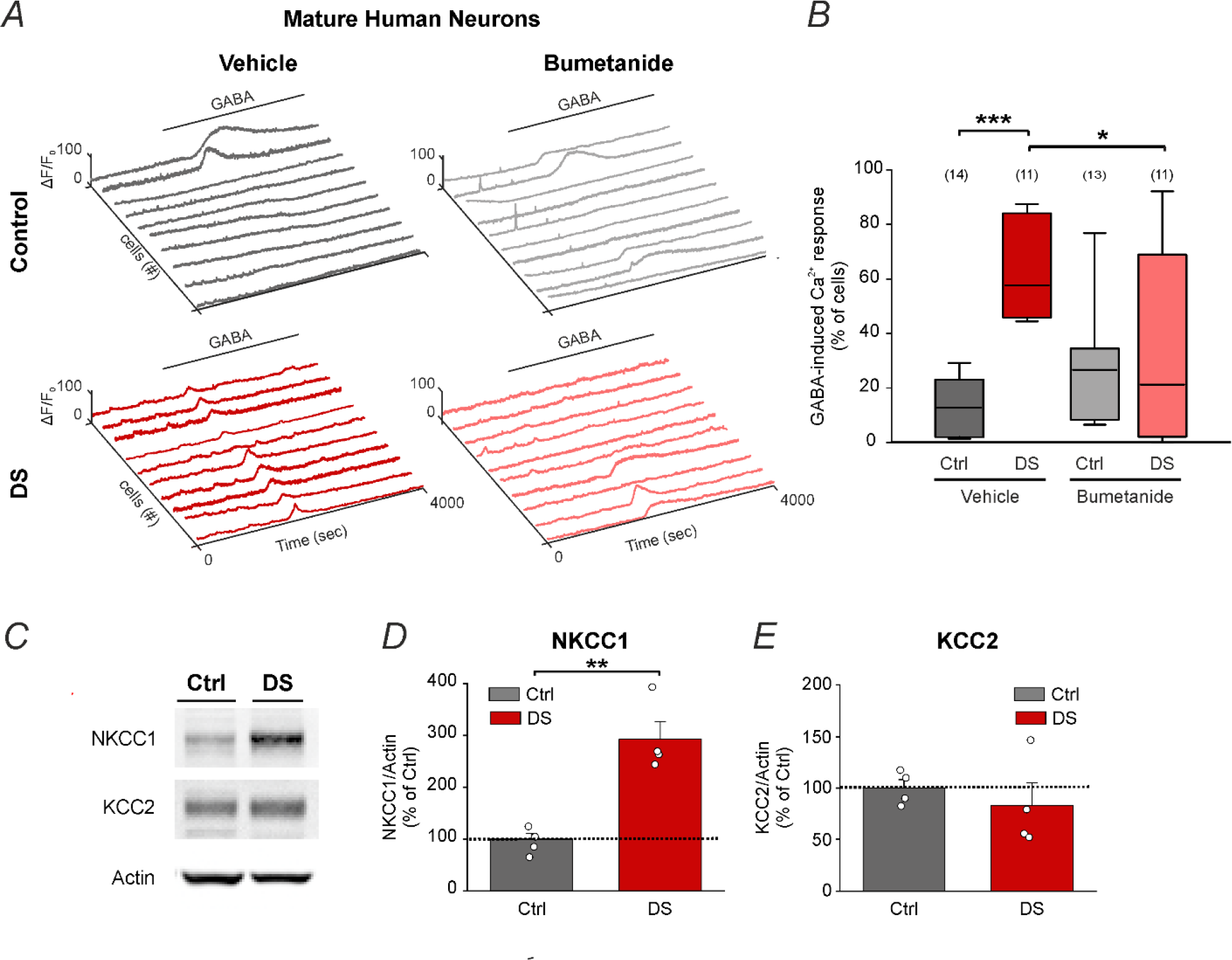
Mixed subpopulations of neurons with depolarizing or hyperpolarizing GABA signalling are present in human isogenic control and trisomic iPSC-derived neurons obtained from a person with DS. **A)** We analysed trisomic neurons and corresponding isogenic control (euploid) neurons derived from iPSCs obtained from an individual with a mosaic form of DS. Representative calcium traces of iPSC-derived control and trisomic (DS) neurons pretreated with vehicle (0.01% DMSO) or the NKCC1 inhibitor bumetanide (10 μM) upon bath application of GABA (100 μM) at 60 days following plating for final differentiation. **B)** Quantification of the percentage of neurons showing depolarizing GABA responses in experiments as in A. Boxplots indicate median and 25th-75th percentiles, and whiskers represent the 5th- 95th percentiles. The numbers in parentheses indicate the number of analysed coverslips for each experimental group (obtained from 5 independent neuronal differentiation experiments). *P<0.05, ***P<0.001, Tukey *post hoc* test following two-way ANOVA. **C)** Representative immunoblots for NKCC1 and KCC2 protein extracts from control and trisomic (DS) neurons. Actin was used as an internal standard. Full blots are shown in Supplementary Fig. 10A. **D)** Quantification of average NKCC1 protein levels (± SEM; expressed as the percentage of control neurons, dotted line) in same experiments as in C. Actin was used as an internal standard. Dots indicate values of individual coverslips (obtained from 2 neuronal independent differentiation experiments). **P<0.01, Student’s t test. **E)** Quantification of average KCC2 protein levels (± SEM) in the same experiments described in D. Dots indicate values of individual coverslips (obtained from 2 neuronal independent differentiation experiments).

Overall, these results demonstrated the presence of a subpopulation of neurons presenting depolarizing GABAA signalling in both human control and DS mature cultures, with DS cultures presenting a lager NKCC1-dependent subpopulation.

### Some evidence of heterogeneous subpopulations of WT mature neurons with hyperpolarizing or depolarizing GABAAR-mediated responses has been hinted in the literature

Since we found clear evidence of heterogeneous subpopulations of neurons with respect to GABAAR- mediated depolarizing or hyperpolarizing responses at increasing scales of network complexity even at mature stages of development in WT animals and in human iPSC-derived neurons, we wondered whether the current literature would support any even course evidence of the same phenomenon. To this aim, we screened the current literature to find publications on the identification of a mixed population of hyperpolarizing or depolarizing GABA-responding WT neurons. Since no publication specifically addressed this issue, we considered the very few publications quickly describing depolarizing GABA responses in the results section and those even simply displaying overlooked data in their figures (Table 1).

**Table 1.**
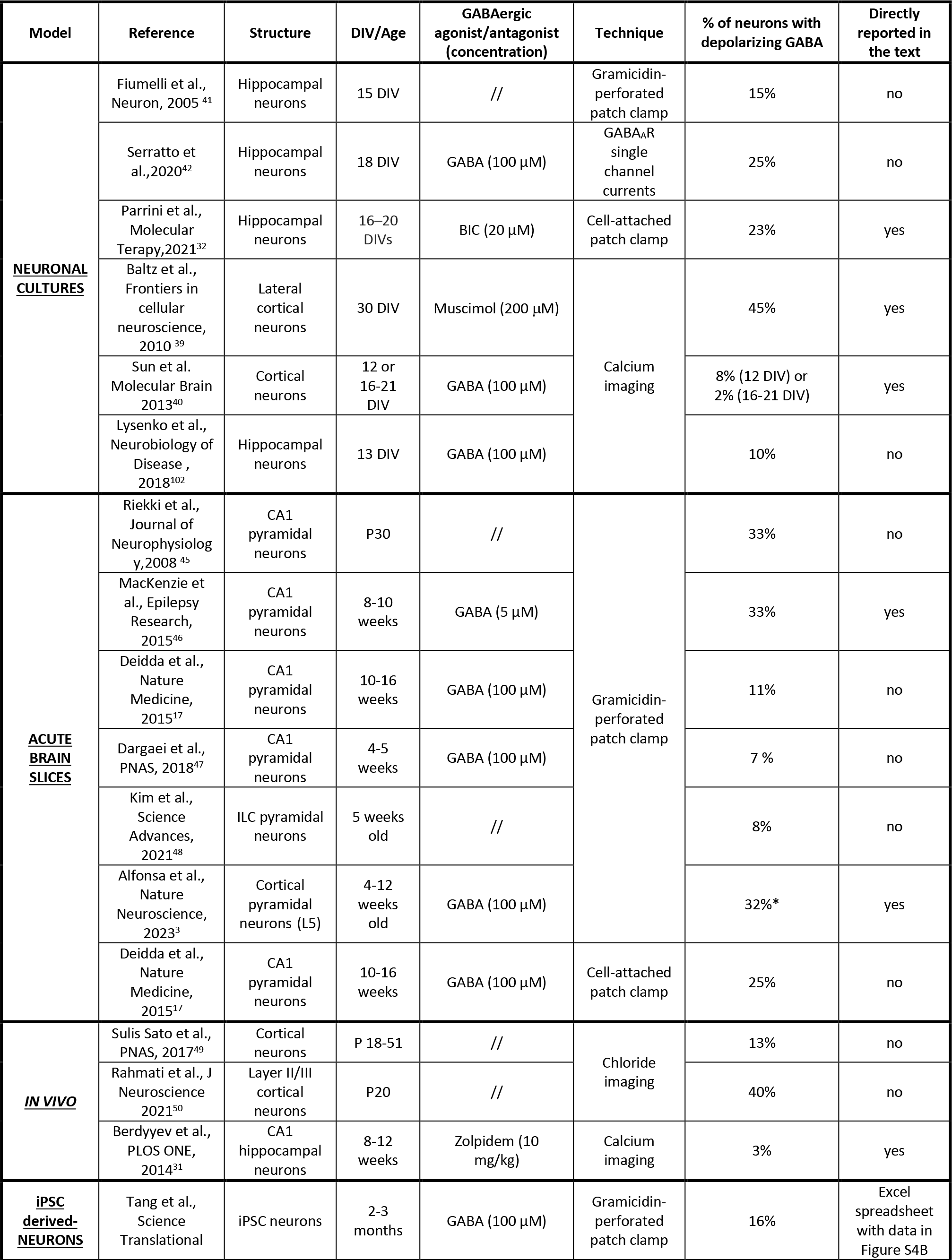
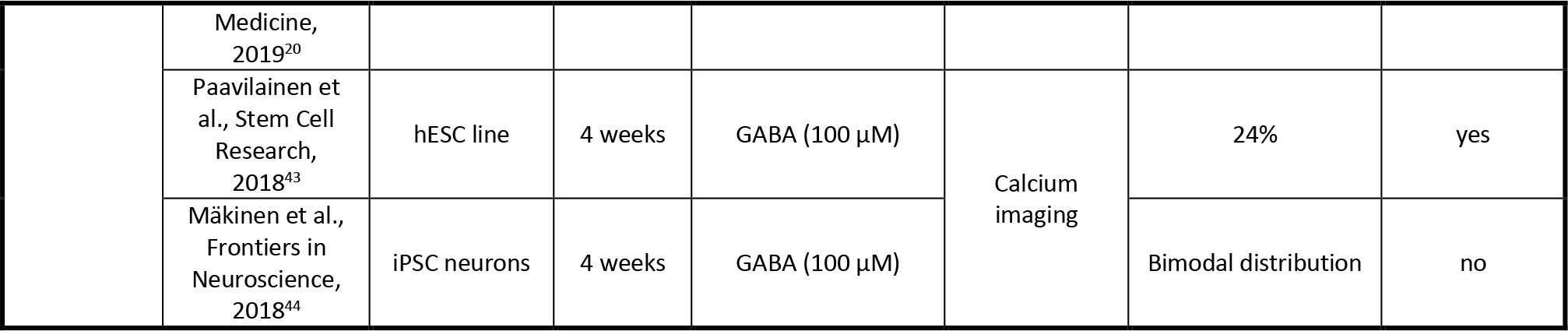
A mixed population of GABA-responding neurons at an increasing scale of complexity has already been described in the literature. Abbreviations: RMP, resting membrane potential, reference value of -65 mV; DIV, days *in vitro*; P, postnatal day; iPSC, induced pluripotent stem cells; hESC, human embryonic stem cells; ILC, infralimbic cortex. *Note: 5% with action potential after GABA puffing (100 μM).

To classify cells as characterized by hyperpolarizing or depolarizing GABA, we relied primarily on what authors explicitly reported in (Ca^2+^ or Cl^-^) imaging and electrophysiological (gramicidin-perforated, GABAAR single channel currents or cell-attached patch-clamp) datasets obtained in rodent neuronal primary cultures, brain slices and *in vivo* experiments, as well as in human neurons derived from control iPSC lines (Table 1). When the classification of cells in depolarizing and hyperpolarizing GABA was not explicitly reported by the authors, we looked at the single-neuron values directly in the figures, and we calculated the approximate percentage of neurons that fell in the two different categories (*i.e.*, hyperpolarizing *vs.* depolarizing GABA) ourselves (Table 1). In particular, for Ca^2+^ imaging experiments, we considered an increase in Ca^2+^ signalling following GABAAR agonist treatments as a depolarizing GABA response. For Cl^-^ imaging, we considered neurons with depolarizing GABA those presenting a high intracellular Cl^-^ concentration (> 40 mM, as in neurodevelopment), which would likely lead to an outward (depolarizing) flow of Cl^-^ through GABAARs. For electrophysiological gramicidin-perforated or GABAAR single channel currents, we considered depolarizing GABA neurons to be those that displayed EGABA higher than the cell RMP. For cell-attached patch-clamp recordings, we considered neurons with depolarizing GABA those showing an increase/decrease in the frequency of action potentials in response to a GABAA receptor agonist/antagonist, respectively.

Overall, we could only find 29 studies that showed data relevant to our research in some way (Table 1 and Supplementary Table 2). In particular, in rodent WT primary neuronal cultures, both Ca^2+^ imaging experiments and gramicidin-perforated patch clamp recordings in the literature reported a percentage of neurons with depolarizing responses to GABAAR stimulation even at mature (15-30 DIV) stages of development (Ca^2+^ imaging: 2-45% of GABA depolarizing neurons ^39, 40^; gramicidin-perforated patch-clamp recordings: 15% of GABA depolarizing neurons ^41^; GABAAR single channel currents: 25% ^42^; cell-attached patch clamp recordings: 23% ^32^ of GABA depolarizing neurons). Notably, such findings were replicated also in mature (4-12 weeks of differentiation) iPSC-derived human neurons (Ca^2+^ imaging: 24% of GABA depolarizing neurons ^43, 44^; gramicidin perforated patch-clamp recordings: 16% of GABA depolarizing neurons ^20^). Interestingly, such heterogeneity in the polarity of GABAAR-mediated responses was not limited to neurons in culture, which may indeed never reach a full and complete maturational stage, but also in acute brain slices from adult mice (gramicidin-perforated patch-clamp: 7-33% ^3, 17, 45–48^; cell-attached patch clamp recordings: 25% ^17^. Most notably, *in vivo* evidence also pointed to a subpopulation of neurons with depolarizing GABAergic responses in juvenile animals after the GABA developmental polarity switch and in fully adult animals (overall age range P18-51; chloride imaging: 13-40% of neurons ^49, 50^; Ca^2+^ imaging: 3% of neurons ^31^; Supplementary Table 1). Moreover, we note here that in further 4 articles of the total 29 articles that we found relevant to our analysis of the literature authors found neurons with depolarizing GABA signalling in some experimental settings but not in others, while in other 5 publications they did not find (*i.e.*, actively reported or just showed in the figures) any subpopulation of mature neurons with depolarizing responses to GABA (Supplementary Table 2). Finally, in 6 articles authors reported only average data values, and we could not use them for our analysis (Supplementary Table 2).

Thus, although the current literature is extremely scant on the topic, it confirmed the presence of an overlooked heterogeneous population of neurons with respect to the polarity and direction of GABAAR- mediated signalling in human and murine WT mature cells *in vitro* and for mouse neurons also *in vivo*.

## DISCUSSION

The current mainstream view regarding the regulation of mature neuronal network activity in the brain suggests that a small number of inhibitory GABAergic neurons dampens the firing of the large majority of excitatory glutamatergic neurons by hyperpolarizing postsynaptic cells. However, in immature neurons, GABAergic signalling is actually depolarizing and mostly excitatory. The polarity of GABAergic signalling switches to inhibitory during brain development due to the decrease in [Cl^-^]i by upregulation of the expression of the Cl^-^ exporter KCC2 ^15^. Interestingly, experimental and computational simulations have nevertheless shown that even in mature neurons with already fully inhibitory GABAAR-mediated signalling, small changes in EGABA can potentially shift GABA inhibition into excitation ^9, 51^. Therefore, we reasoned that it is possible that the GABA polarity switch may never be fully accomplished in the entire population of neurons or that some neurons may keep oscillating between depolarizing and hyperpolarizing GABA signalling even when they become fully mature. Here, we experimentally indicated that a small subpopulation of WT mature neurons showed high [Cl^-^]i and excitatory GABAergic signalling and that this subpopulation was intermingled with the larger majority of neurons with inhibitory GABA signalling both *in vitro* and *in vivo*. This mixed population of GABA-responding neurons started to appear during development when the proportion of neurons with hyperpolarizing *vs.* depolarizing GABAergic signalling increased progressively, yet without reaching wholeness. However, only tracking of each individual neuron over time will reveal whether the population of mature neurons with depolarizing GABAAR signalling is derived from neurons that never fully mature or neurons with peculiar proprieties in regulating [Cl^-^]i. In fact, (at least) some neurons appear to have significant cellular microdomains of diverse intracellular Cl^-^ concentration (*i.e.*, mosaicism of GABAergic signalling in the axon and axonal initial segment, in the soma, and along dendrites). While this heterogeneity of GABAAR-mediated responses at the level of cellular compartments is well accepted ^52–56^, the presence of a heterogeneous subpopulation of neurons with depolarizing *vs.* hyperpolarizing GABA responses in mature neuronal networks has been, at times, hinted by some scant literature (Table 1), but it is not yet a clear and well-established concept in the neuroscience field. This is in spite of the fact that GABA modulation of brain activity in adult neuronal networks is among the most studied topics in neuroscience.

Our data and careful analysis of the current literature (Table 1) in fact clearly point to the presence of a mixed population of neurons with hyperpolarizing and inhibitory or depolarizing and excitatory GABAergic signalling in mature networks. On the other hand, our review of the literature also highlighted that some authors did not report the same heterogeneity in GABAAR-mediated responses (Supplementary Table 2). This may simply be due to the different experimental conditions or analysis. It is plausible that in certain experimental designs permissive of a small number of recorded cells, the very low percentage of cells with depolarizing GABA responses may have been missed or may have been interpreted as outliers in the statistical analysis.

Our results were strengthened by the fact that we observed the presence of neurons with mixed responses to GABA signalling at three diverse layers of neuronal network complexity ranging from *in vitro* (primary mouse cultures and human iPSC-derived neurons) to *ex vivo* (mouse acute brain slices) and *in vivo* (freely moving mice). Interestingly, the size of the population of mature neurons with depolarizing and excitatory GABA responses increased from cell cultures to brain slices. On the other hand, we found a smaller number of GABA-depolarizing neurons *in vivo* than what we observed at other levels of complexity. This outcome might be explained in part by the fact that to unmask the population of neurons with excitatory GABA responses, we used diverse pharmacological approaches (GABA and bicuculline application *in vitro* and diazepam application *in vivo*) or due to the heavy pharmacokinetic and pharmacodynamic processing *in vivo* compared to *in vitro*. Moreover, whereas GABA directly activates GABAARs causing complete suppression of firing activity in WT neurons, diazepam is only a positive modulator of GABAARs, thus leading to a smaller reduction in neuronal activity ^57, 58^, as revealed also by our own data in cell cultures. Moreover, *in vitro* culturing or brain slicing procedures may affect chloride homeostasis *per se* ^59^. Finally, it is also possible that the very high number of cells that were nonresponsive to diazepam *in vivo* are indeed among the cells with high levels of NKCC1 expression, as their responsiveness to bumetanide would indicate in our experiments. Anyhow, our *in vivo* results are perfectly in line with previous reports indicating a small percentage of neurons with increased [Cl^−^]i ^49^ or depolarizing responses to another GABAAR-positive allosteric modulator (Zolpidem) in freely behaving animals ^31^.

Both experimental and computational studies have shown that in mature neurons, EGABA value is only a few mV more negative than the RMP; therefore, even small deviations from the set-point in one or both of these measures may cause a switch in the polarity of GABAergic signalling from inhibition to excitation or *vice versa* ^9^. In this framework, it is not surprising that many brain disorders are characterized by depolarizing GABAergic signalling ^14, 16, 26, 60^. In light of our study, it is tempting to speculate that this concept may simply reflect an increase in the percentage number of cells with high [Cl^-^]i (and thus depolarizing and excitatory GABAergic signalling) and not a generalized small increase in the level of [Cl^-^]i in all neurons. Accordingly, we found a large population of mature neurons with excitatory GABAergic responses in the Ts65Dn mouse model of DS at all levels of investigation and in line with the current literature on altered average Cl^-^ homeostasis in adult trisomic mice ^17, 32, 35, 61^.

Although we found that the number of depolarizing neurons represents a small minority compared to the number of hyperpolarizing ones in WT and even in Ts65Dn mature brains, this finding may still have important functional implications. However, thus far, manipulations of this population of neurons with depolarizing GABA in adult WT animals by pharmacological treatment with NKCC1 inhibitors ^14, 17, 35, 47, 61–64^, genetic deletion of NKCC1 or its downregulation by RNA interference ^32, 65–68^ have been shown not to significantly affect cognitive (novel object recognition test, Y maze, object location test, conditional fear conditioning test, and operant reversal learning test ^17, 32, 35, 47, 48^), social (three chamber and male– female interaction^35^) and motor functions (tail suspension, motor coordination, and forced swim test ^69, 67, 70, 71^) or to significantly increase anxiety (open field test and elevated plus maze ^69^). Further experiments addressing more subtle and specific behaviours may reveal a role for the population of neurons with depolarizing GABA responses in WT animal behaviour under physiological conditions. Anyhow, modulating NKCC1 with pharmacological inhibitors or RNA interference in mouse models of a number of pathologies (including DS) as well as in patients has shown positive behavioural outcomes ^14, 26, 72^. This finding indicates that the small population of neurons with depolarizing GABA responses may have in fact an important role in pathological conditions. Our *in vivo* experiments on behaving mice and computational analysis add substantial knowledge to this mostly *in vitro* experiment-based broad discussion in the field aimed at directly addressing whether the population of neurons with depolarizing GABA signalling have a pathological role in the many mouse models of brain disorders characterized by depolarizing GABA signalling. We found here that adult DS mice had a larger population of depolarizing-GABA neurons than WT mice *in vivo*. As a consequence, enhancing GABAergic signalling by diazepam treatment in adult DS mice led to a paradoxical anxiety increase in the dark-light behavioural test, which was rescued by bumetanide pretreatment. Since bumetanide treatment significantly reduced the number of mature neurons with excitatory GABA responses in Ts65Dn neurons in cultures, brain slices and *in vivo*, the rescue of paradoxical responses to benzodiazepines in Ts65Dn animals by bumetanide treatment indeed points to a pathological role of the population of neurons with depolarizing GABA in DS. On the other hand, and in agreement with the current literature ^36^, we found a substantial lack of anxiolytic diazepam activity, but without a clear paradoxical response in Ts65Dn mice in the open field test and elevated plus maze, possibly indicating a low sensitivity of these tests in revealing this phenotype. Despite this, we found that bumetanide treatment strongly restored anxiolytic diazepam action in adult Ts65Dn mice in both tests.

Although benzodiazepines are used as effective anxiolytics and sedatives worldwide for a vast number of therapeutic indications, adverse paradoxical reactions (*i.e.*, anxiogenic *vs.* anxiolytic effects) have been described in subjects with neurodevelopmental disorders (*e.g.,* autism and schizophrenia) ^73–75^. Moreover, paradoxical reactions to benzodiazepines, including increased emotional release, excitement, excessive movement, panic attacks and even hostility and rage, were reported in subjects with several predisposing risk factors, such as young age, advanced age, genetic predisposition (*i.e.,* diabetes mellitus and atrial fibrillation), alcoholism, and psychiatric and/or personality disorders (*i.e.,* depressive bipolar disease, bipolar affective disorder, borderline personality disorder, manic depression and chronic insomnia ^74, 75^). This may indicate that an increased population of neurons with excitatory GABA responses may also be present in these pathologies, as we described here in adult DS animals. Moreover, our results on the modulation of the GABAAR-dependent effect of diazepam (both on anxiety behaviour and Ca^2+^ imaging *in vivo*) by systemic bumetanide pretreatment indicate that NKCC1 has a role in mediating

GABAergic responses in the brain *in vivo*. This finding is very relevant, considering the poor blood-brain barrier penetration of bumetanide, which at times has cast doubts on its Cl-dependent (and thus GABA-dependent) mechanism of action ^76, 77, 78, 79^.

To increase the translational value of our data, we validated our results in human mature neurons derived from a control iPSC line and a trisomic iPSC line both of which obtained from the same person with mosaic DS. These data strongly support the design of current (EudraCT 2015-005780-16) and future clinical trials with NKCC1 inhibitors in DS. Moreover, the increased expression of NKCC1 corresponding to an increased population of trisomic cells with depolarizing GABA (which was inhibited by bumetanide) suggests that iPSC-derived neurons from patients could be used to test NKCC1 levels as a biomarker of depolarizing GABAergic activity or even test responsiveness to pharmacological treatments of particular individuals with DS. Thus, neurons derived from patient iPSC lines could be used for patient stratification in future clinical trials for DS and possibly for many other brain disorders characterized by impaired chloride homeostasis ^14^.

In conclusion, we demonstrated the presence of a mixed population of neurons responding to GABA with hyperpolarizing/inhibitory or depolarizing/excitatory actions in mature human and murine neuronal networks at various levels of complexity. While the physiological function of the population of neurons with depolarizing/excitatory GABA-driven responses in mature WT animals needs further investigation, the increase in this population in disease may play a pivotal role in determining key aspects of the pathology. As the list of brain disorders associated with depolarizing GABA signalling is already very large and increases every day, our experiments on a mouse model of one of these pathologies put forward a rather simple concept. Brain networks can normally tolerate (and possibly need) a small heterogeneous subpopulation of neurons with depolarizing GABA signalling, but if disease-associated changes impinge on the subtle balance between the extent of subpopulations of neurons with excitatory *vs.* inhibitory GABAergic signalling, very negative consequences can arise.

## METHODS

### Animals

The Ts65Dn colony 80 was maintained by crossing Ts65Dn females to C57BL/6JEi x C3SnHeSnJ (B6EiC3) F1 males (Jackson Laboratories). Dp(16)1Yey/+ 81 males were purchased from The Jackson Laboratory and used to create a colony by mating with C57BL6/J females (Charles River). All animals were genotyped by PCR as previously described 81, 82. A veterinarian was employed to monitor health and comfort of the animals. Mice were housed in a temperature-controlled room with a 12:12 hour dark/light cycle and ad libitum access to water and food. All animal procedures were approved by IIT licensing, the Italian Ministry of Health (D.Lgs 26/2014) and EU guidelines (Directive 2010/63/EU).

### Ethical approval declaration

All animal experiments were performed in accordance with the guidelines established by the European Community Council Directive 2010/63/EU of September 22nd 2010 and were approved by the Italian Ministry of Health (authorizations no: 829/2015-PR and 658/2016-PR to A.C.).

Primary neuronal culture. Primary neuronal cultures were prepared from WT and Ts65Dn mouse pups at postnatal day 2 (P2), as previously described ^18, 35, 83^. In brief, brains were dissected under a stereomicroscope in ice-cold buffer (DB) composed of Hank’s balanced salt solution (HBSS; Gibco) supplemented with 6 mg/mL glucose, 3 mg/mL bovine serum albumin (BSA), 5.5 mM MgSO4, 5 g/mL gentamycin and 10 mM Hepes, pH 7.4 (all from Sigma). Cortical or hippocampal tissue was minced and then enzymatically digested with 0.25% trypsin in HBSS containing 0.6 mg/mL deoxyribonuclease I (DNAse; Sigma) for 5 min at 37°C. Tissue chunks were washed in DB, incubated for 5 min in DB supplemented with 1 mg/mL of Soybean trypsin inhibitor (Sigma) and mechanically dissociated in DB supplemented with 0.6 mg/mL DNAse. Cells were passed through a 40 m strainer and then centrifuged (110 x g for 7 min at 4°C) to remove cellular debris. Cells were plated on glass coverslips (Menzel Gläser), 6-well plates (BD Falcon) or Microelectrodes Arrays (MEAs, Multichannel System, Germany) all coated with poly-L-lysine (Sigma; 0.1 mg/mL in 100 mM borate buffer, pH 8.5) at a density of 250-500 cells/mm2. Neurons were maintained in a culture medium consisting of Neurobasal-A supplemented with 2% B-27, 1% GlutaMax and 5 g/mL gentamycin (all from Gibco) at 37 °C in humidified atmosphere (95% air, 5% CO2).

Human iPSC-derived neurons. The previously generated Down syndrome cell line DS4 of human induced pluripotent stem cells (iPSCs) and the corresponding isogenic control line DS2U (Weick et al., 2013) were obtained from WiCell (Madison, Wi). Both iPSC lines were obtained from an individual with a mosaic form of DS. Human DS and control iPSCs were cultured and maintained in mTeSR1 medium (Stem Cell Technologies) in tissue culture dishes coated with hESC-qualified Matrigel (Corning). Cells were routinely passaged as aggregates with ReLeSR (Stem Cell Technologies) dissociation reagent.

For neuronal differentiation, a commercially available differentiation method (Stem Cell Technologies) based on the generation of embryoid bodies (EBs) was employed to first generate neural progenitor cells (NPCs). In brief, EBs were generated by seeding single-cell suspension of iPSC in AggreWell-800 plates (Stem Cell Technologies) in STEMdiff Neural Induction Medium (NIM; both from Stem Cell Technologies) in the presence of the ROCK inhibitor Y-27632 (10 M). After 5 days, EBs were removed and plated onto Matrigel-coated dishes (BD Falcon) in NIM, where they attached and formed neural rosettes. After further 5 days, the rosettes were isolated with Neural Rosette Selection Reagent (Stem Cell Technologies) and re- plated onto Matrigel-coated dishes in NIM. After additional 5-7 days, NPCs outgrew from rosette and formed a monolayer of cells. NPCs were then detached with StemPro Accutase (Gibco) and re-plated on Matrigel-coated dishes in NPC Medium (DMEM/F12, 2% B27, 1% N2, 1% Pen-strep; all from Gibco) supplemented with 20 ng/mL bFGF (Peprotech). Cells were passaged using StemPro Accutase and re- plated at 72,000 cells/cm2 in complete NPC Medium for maintenance.

For neuronal differentiation, NPCs were first pre-differentiated by plating into 10 cm plastic dishes coated with Matrigel in complete NPC Medium as above. The next day, a complete media change with NPC Medium was performed. On the second day, half volume of Neuronal Differentiation Medium (NDM; Neurobasal A, 2% B27, 1% N2, 1% Glutamax, 1% Pen-strep, 20 ng/mL BDNF and GDNF, 100 M dibutyryl- cAMP and 200 nM Ascorbate) was added to the culture. On the fourth day, half volume of the media was changed with fresh NDM media. On the fifth day, cells were detached with StemPro Accutase, counted in a haemocytometer chamber, and re-plated for final differentiation at the density of 18,000 cells/cm2 on PLO/Laminin coated glass coverslips (Menzel Gläser). Coverslips were first coated overnight with poly-L-ornithine (Sigma; 0.05 mg/mL in 100 mM borate buffer, pH 8.5) in the incubator at 37oC. On the day of plating, coverslips were washed with autoclaved milliQ water and left to dry before coating with 1 g/mL of Laminin (Sigma) diluted in DMEM/F12 for 4 hours in the incubator at 37oC. Laminin was removed just before plating the cells. For culture maintenance, half of the medium was changed three times a week with the complete NDM medium supplemented with 1 μg/mL of Laminin. Cells were used for Ca2+ imaging or lysed for biochemistry 60 days after plating for the final differentiation.

In vitro calcium imaging. Mouse primary neuronal cultures. Calcium imaging experiments on WT or Ts65Dn hippocampal primary neuronal cultures grown on glass coverslips were performed with the Ca2+- sensitive dye Fluo4-AM (Molecular Probes). Coverslips at different days in vitro (2, 7, 15 or 21) were loaded with 2.5 M Fluo4-AM in extracellular solution (NaCl 145 mM, KCl 5 mM, CaCl2 2 mM, MgCl2 1 mM, HEPES 10 mM, D-glucose 5.5 mM, pH 7.4) for 20 minutes in the dark at room temperature. Coverslips were then transferred to a holding chamber and perfused (2 mL/min) with the same extracellular solution for 2-3 minutes before imaging. Live imaging was performed with a Swept Field Confocal (SFC) microscope (Nikon) equipped with a 20X air-objective (NA 0.75) under constant perfusion. After 10-15 minutes of baseline imaging, neurons were bath stimulated with 100 M GABA or 10 M muscimol (both from Sigma) for additional 8-10 minutes. Following GABA or muscimol washout, neurons were stimulated with 30 mM KCl to depolarize cells and evaluate the total number of viable neurons present in the field of view. In some experiments, we used the GABAAR antagonist bicuculline (Sigma; 100 M), the GABABR antagonist CGP55845 (Tocris; 10 M) or the L-type voltage-gated calcium channel blocker nifedipine (Tocris; 10 M) in combination to GABA stimulation (100 M). For these experiments, we first recorded 10-15 minutes of baseline (as above), then we performed 10-15 minutes of imaging in the presence of the tested drug, followed by 8-10 minutes of imaging in the presence of GABA (100 M) plus the tested drug. Finally, after washout of the drugs, neurons were depolarized with 30 mM KCl (as above). In other experiments, neurons were pretreated with 10 M of bumetanide (Sigma) or the corresponding vehicle (0.01% DMSO) during Fluo4-AM loading and subsequently imaged (as above), but under constant perfusion with extracellular solution containing either bumetanide or DMSO. One field of view for each coverslip was acquired. iPSC-derived human neurons. For calcium imaging experiments on control and DS iPSC-derived human neurons, cultures were grown on glass coverslips for 60 days after final plating. Cells were loaded with 2.5 M of Fluo4-AM directly in their culture medium for 20 minutes in the incubator. Coverslips were then transferred to a 35 mm dish containing an imaging solution (NaCl 95 mM, KCl 5 mM, CaCl2 1.8 mM, MgCl2 0.8 mM, NaH2PO4 1 mM, NaHCO3 26 mM, HEPES 10 mM, D-glucose 10 mM, pH 7.4) before being transferred to a holding chamber and perfused (2 mL/min) with the same solution for additional 2-3 minutes. Similar to WT and Ts65Dn mouse primary neurons, human iPSC-derived neurons were pretreated with 10 M bumetanide (Sigma) or the corresponding vehicle (0.01% DMSO) during Fluo4-AM loading and subsequently imaged with constant perfusion with imaging solution containing either bumetanide or DMSO. Live imaging was performed with an inverted microscope (Nikon) equipped with a 20X air-objective (NA 0.75), a pE-300 LED illumination system (CoolLED) and an iXON EMCCD camera (Andor Technology). After 10 minutes of baseline imaging under constant perfusion, neurons were bath stimulated with 100 M GABA (Sigma) for additional 8 minutes. Following GABA washout for 4 minutes, neurons were stimulated with 30 mM KCl to depolarize cells and evaluate the total number of viable neurons present in the field of view. One field of view for each coverslip was acquired.

For all cells, image analysis was performed by NIS-Elements software (Nikon). First, regions of interest (ROIs) were placed on the cell body of individual neurons identified in video frames, following KCl depolarization of cells (representing the maximum fluorescence of viable neurons). Next, average Fluo4 fluorescent intensity for each ROI was measured by NIS-Elements software on every frame of the recording to generate the corresponding calcium traces for each cell. The raw calcium traces obtained for each neuron were then transformed to relative changes in fluorescence (F/F0) over time. The value of F/F0 for each cell was calculated as (F-F0)/F0, where F0 is the minimal fluorescence signal for a given cell. An operator blinded to the experimental sample code analyzed the calcium traces and scored neurons showing GABA-dependent depolarizing responses. Neurons were scored positive if GABA or muscimol application elicited a calcium transient that was at least 50% higher than the basal fluorescence. The percentage of neurons showing GABA-dependent depolarizing responses was calculated for each field of view over the total number of viable neurons (i.e., neurons depolarized by KCl application) in the same field of view.

In vitro chloride imaging. Imaging of intracellular Cl- in WT or Ts65Dn primary hippocampal neuronal cultures grown on coverslips was performed with the fluorescent chloride-sensitive indicator MQAE [N- (Ethoxycarbonylmethyl)-6-Methoxyquinolinium-Bromide; Molecular Probes], as previously described ^18, 35^. The MQAE dye detects Cl- ions via diffusion-limited collisional quenching, resulting in a concentration-dependent decrease of fluorescence emission following an increase in Cl- concentration 84. Therefore, a decrease in MQAE fluorescence is indicative of a higher [Cl-]i and vice versa. Hippocampal neurons at 15 DIV were loaded with 5 mM MQAE in their culture medium for 30 minutes at 37°C in the presence of bumetanide (10 M) or the corresponding vehicle (0.01% DMSO). Coverslips were then transferred to a holding chamber and perfused (2 mL/min) with extracellular solution (as above) containing either bumetanide or vehicle for 5 minutes before imaging. Images were acquired with a Nikon A1 scanning confocal microscope equipped with a 20X air-objective (NA 0.75). MQAE was excited with a 405 nm diode laser and fluorescence collected with a 525/50 nm band-pass emission filter. All excitation and acquisition parameters (laser intensity, PMT offset and gain) were keep constant throughout experiments. Image analysis was performed with NIS-Elements software (Nikon) by measuring the mean fluorescent intensity of ROIs placed on the cell body of individual neurons from 6 randomly selected fields for each coverslip. For each experiment, the average fluorescent intensity of all ROIs from a coverslip was normalized to the average fluorescent intensity of control samples (WT neurons treated with vehicle) in the same experiment. Pseudo-color images were generated with ImageJ software (http://rsbweb.nih.gov/ij/). The beanplots were generated with R package (http://cran.r-project.org/web/packages/beanplot/index.html).

Biochemistry. Primary neuronal cultures, iPSC-derived neurons or brain samples from WT, Ts65Dn and Dp(16) mice were lysed in ice-cold RIPA buffer (1% NP40, 0.5% Deoxycholic acid, 0.1% SDS, 150 mM NaCl, 1 mM EDTA, 50 mM Tris, pH 7.4) containing 1 mM PMSF, 10 mM NaF, 2 mM sodium orthovanadate and 1% (v/v) protease and phosphatase inhibitor cocktails (Sigma). Samples were clarified through centrifugation at 20,000 x g at 4°C, and the protein concentration was determined using the BCA kit (Pierce). For immunoblot analysis, protein extracts were prepared in lithium-dodecyl-sulfate (LDS) sample buffer (ThermoFisher Scientific) containing 50 mM dithiothreitol (DTT). To avoid NKCC1 or KCC2 protein aggregation/precipitation, samples were not heat-treated before loading 85, 86. Equal amounts of proteins were run on 4-12% Bis-Tris, NuPAGE gels (ThermoFisher Scientific) with MOPS buffer and transferred overnight at 4°C onto nitrocellulose membranes (GE Healthcare) with Tris-Glycine transfer buffer (25 mM Tris-base, 192 mM glycine, 20% methanol). Membranes were probed with mouse anti- NKCC1 (clone T4c, Developmental Studies Hybridoma Bank; 1:4,000), rabbit anti-KCC2 (Millipore, catalog no. 07-432; 1:4,000), and rabbit anti-Actin (Sigma, catalog no: A2066; 1:10,000), followed by HRP-conjugated goat anti-mouse, goat anti-rabbit secondary antibodies (ThermoFisher Scientific; 1:10,000). Chemiluminescent signals were revealed with SuperSignal West Pico substrate (Pierce) and digitally acquired on a LAS 4000 Mini imaging system (GE Healthcare). Alternatively, a fluorescent Cy5-conjugated goat anti rabbit secondary antibody (GE Healthcare; 1:5,000) was used and the fluorescent signal acquired with a Typhoon fluorescent scanner (GE Healthcare). Band intensities were quantified using ImageQuant software (GE Healthcare). For some experiments, membranes were first probed with anti-NKCC1 antibody, stripped and then re-probed with anti-KCC2 antibody. Specificity of the anti-NKCC1 or anti-KCC2 antibodies was previously verified on brain samples from NKCC1 and KCC2 deficient mice 87.

### MEA recordings

Primary neuronal cultures. WT and Ts65Dn primary hippocampal neuronal cultures were grown on MEAs consisting of 60 TiN/SiN planar, round electrodes (30 μm diameter; 200 μm inter-electrode distance) divided into six separated wells (Multichannel Systems). Each well contained nine recording electrodes, arranged in a 3 × 3-square grid, and one big ground electrode. The network activity of all neuronal cultures was recorded by means of the MEA60 System (Multichannel Systems). After 1200 × amplification, signals were sampled at 10 kHz and recorded using a commercial software (MC-Rack software, Multichannel Systems). To reduce thermal stress of the neurons during each experiment, cultures were recorded in their original medium, kept at 37 °C by means of a controlled thermostat (Multichannel Systems) and covered by a Polydimethylsiloxane (PDMS) cap to avoid evaporation and prevent changes in osmolarity ^32, 88^. Additionally, a custom incubation chamber was used to maintain a controlled humidified atmosphere consisting of 95% air and 5% CO2 during the entire recording, as previously reported^32, 88, 89^.

Cultures at 20–21 DIVs were pretreated with bumetanide (10 uM) or with the corresponding vehicle (0.01% DMSO) for 30 minutes in the incubator prior to each recording session, and then accommodated on the MEA set-up for 15 min, to let the culture adapt to the new environment and reach a stable level of activity^32, 88^. Next, basal spontaneous network activity was recorded for 30 min. After recording basal activity, bicuculline (20 μM; Sigma), GABA (100 μM; Sigma) or diazepam (1 μM; Tocris) were added by directly pipetting in the medium, and activity was recorded for an additional 40 min. Since we previously noticed that mechanical perturbation from the pipette injection in the medium could cause a temporary instability in the firing rate^32, 89^, we discarded the first 10 min at the beginning of the 40-min recording phase.

Data analysis was performed offline with MATLAB software (MathWorks) using the custom software package SPYCODE90. The analysis was performed as previously described^32, 88^. Briefly, raw traces were high-pass filtered (>300 Hz) to isolate spike events from the low fluctuation of the signal (LFP). Spike detection was computed offline using a custom software, to discriminate spike events from the noise (Bologna et al. 2010). To characterize the level of the activity of the recorded cultures, the mean firing rate (MFR) for each MEA was defined as the mean number of spikes per second (spikes/sec) over the total recording time and computed. Active electrodes were defined as those showing a firing rate greater than 0.02 spikes/s. This low threshold guaranteed that only electrodes that were not covered by cells or those electrodes recording very few spikes were excluded by the analysis, and that all other electrodes were retained for analysis.

To evaluate the effect of the drug administration on the firing-rate activity, the MFR ratio, defined as the MFR after drug treatment divided by the MFR during baseline (MFRdrug/MFRbaseline) was computed for each active electrode, and then averaged for each culture.

To assess the presence of a mixed population of neurons, statistically significant changes induced by drug treatments (either GABA, bicuculline or diazepam) in comparison to the basal condition were analyzed at the level of the single electrode with the bootstrap method, as previously described^25, 32^. Briefly, the peak train for each time experimental segment (basal or drug) was divided into 1-min bins. The MFRs recorded during each 1-min bin for the two time segments for each well were randomly shuffled into two groups for 10,000 times. The differences between the means of the randomly shuffled groups produced a null distribution. For each electrode, the real difference between the basal and the drug values was computed and assessed whether it fell outside the 95% confidence interval of the null distribution.

### Hippocampal brain slices

Adult (2-3 months old) WT and Ts65Dn male mice were perfused intracardially with ice-cold oxygenated (95% O2 and 5% CO2) artificial cerebrospinal fluid (ACSF; composed of 115 mM NaCl, 1.25 mM NaH2PO4, 26 mM NaHCO3, 3.5 mM KCl, 1.3 mM MgCl2, 2 mM CaCl2, 25 mM glucose and 1 mM L-Ascorbic Acid, pH 7.4) under deep isoflurane anesthesia and then decapitated. Brains were quickly removed from the skull into a bowl containing ice-cold oxygenated ACSF and then placed into a small beaker containing oxygenated slushy cutting solution (composed of 230 mM sucrose, 3.5 mM KCl, 1.25 mM KH2PO4, 26 mM NaHCO3, 2 mM MgSO4, 0.5 mM CaCl2, 25 mM D-glucose, 1 mM L-Ascorbic Acid, 3 mM Pyruvic Acid, pH 7.4) for 90-120 seconds. Hippocampus-entorhinal cortex slices (400 μm thick) were horizontally cut with a VT1000S vibratome (Leica Microsystem) in the same solution. After separating the two brain hemispheres with a scalpel blade, slices were gently transferred to a beaker by using an inverted Pasteur pipette and rinsed twice with standard oxygenated ACSF. The slices were then transferred to a commercial holding chamber (KF Technologies) containing oxygenated ACSF and incubated for approximately 20 minutes at 32 °C and then for at least 1 hour at room temperature for recovery.

For brain slice recordings on MEA, we substituted the standard recording ring of the MEA with a customized recording chamber inspired by the design of patch-clamp recording chambers; this provided stable and reliable laminar flow that we kept to a rate of 2 mL/min91.

Slices were pre-incubated with oxygenated recording ACSF (composed of 115 mM NaCl, 1.25 mM NaH2PO4, 26 mM NaHCO3, 3.5 mM KCl, 1 mM MgCl2, 2.4 mM CaCl2, 25 mM glucose and 1 mM L-Ascorbic Acid, pH 7.4), containing either vehicle (0.01% DMSO) or bumetanide (10 μM) for 30 minutes, and then transferred to the MEA recording chamber while continuously perfusing with the same solution. Then, slices were let to acclimatize into the MEA setup for 15 minutes to adapt to the new environment and reach a stable level of activity. Next, basal spontaneous network activity was recorded for 30 min. Upon bath application of either bicuculline (20 μM) or GABA (100 μM), additional 15 minutes were waited to ensure complete exchange of the solution in the chamber before recording further 30 min of activity.

Data analysis was performed offline with the MATLAB software (MathWorks) using the custom software package SPYCODE90. Only electrodes positioned in the CA1 region were mapped and analyzed using a custom-made software that allowed the user to select the electrodes that correspond to specific structures of interest of the brain slice91. Raw signals were high-pass filtered (>300 Hz) to isolate the spiking activity. Once spikes were detected using the same algorithms used for cultures, the MFR (spikes/sec) for each slice were computed. Active electrodes were considered those showing a firing rate greater than 0.02 spikes/s, in line with the threshold used for cultures.

To evaluate the effect of the drug administration on the firing activity the MFR ratio, defined as the MFR after drug treatment divided by the MFR during baseline (MFRdrug/MFRbaseline), was computed for each active electrode and then averaged for each slice. In line with the analysis for primary neuronal cultures, differences in the spiking frequency induced by drug treatments at the level of the single electrodes were evaluated to assess the presence of a mixed population of neurons. However, to obtain a reliable null distribution after the bootstrapping procedure, a consistent and well-distributed activity during the recording is required. Therefore, given the more scattered distribution of the firing activity recorded in hippocampal slices compared to primary cultures, a hard threshold of at least 15% change in firing activity was adopted.

Animal treatment and behavioral testing. For behavioral experiments, 3-4 month old male Ts65Dn and WT control littermates were randomly assigned to the diverse experimental groups. Animals were treated daily by intraperitoneal (i.p.) injection for two weeks with either bumetanide (Sigma; 0.2 mg/kg body weight) or vehicle (2% DMSO in saline). On the day of behavioral testing, bumetanide or vehicle injections were given about 1 h before the beginning of the behavioral task followed by diazepam (2 mg/Kg in saline) or saline IP injections 30 minutes before the begging of the test. For these behavioral experiments, injectable diazepam (Valium® 5 mg/mL from Roche) was diluted to 0.2 mg/mL with saline prior to administration, as previously described ^36^.

Animal behavior was video-recorded throughout the experimental sessions by ANY-maze software 92. The operator was blind to both genotype and treatment groups. The open field and elevated plus maze tests were performed in dim light illumination (12-15 Lux), while the dark-light test was conducted under strong illumination (180 Lux). WT and Ts65Dn mice were always evaluated in parallel and within the same time schedule. To minimize olfactory cues from previous trials, all used apparatuses were thoroughly cleaned with 70% ethanol and dried at the beginning of each animal task. Three days before starting the behavioral experiments, mice where handled once a day for 5 min each.

Open field test. This test measures the preference of mice for the central or peripheral area of a square arena. Mice were habituated to the test room for 30 minutes and then placed in the center of a grey acrylic arena (44x44 cm) for 10 minutes of spontaneous exploration. The distance traveled and the time spent in the central or peripheral area of the arena by mice were automatically measured with ANY-maze tracking software. The central area of the arena was defined as a square of 24x24 cm (10 cm away from each wall).

Dark-light test. The apparatus used for the dark-light test consisted of a box divided in two compartments by a panel with a sliding door. The light and dark compartments were made of transparent or black opaque acrylic, respectively. Animals were transferred with a black cage (to avoid exposure to strong illumination before the test) into the behavior room 30 min prior to the beginning of the test. Mice were then individually placed into the dark comportment of the box under dim-light illumination of the room and after 5 seconds (during which the halogen lamp providing the strong illumination was switched on) the sliding door was opened. Mice were allowed to freely explore the two compartments for 10 min. The total number of transitions to and the time spent in the light chamber was analyzed.

### Elevated Plus Maze test

The Elevated Plus Maze apparatus consisted of four arms (30×5 cm) perpendicularly linked to a central platform (5×5 cm). Two opposing arms were open and the other two enclosed by black walls (15 cm high). The platform height from the floor was 40 cm. The platform was placed in the center of a circular tank to prevent mouse escaping in case of fall from the open arms. Mice were habituated to the test room for 30 minutes and then placed in the center of the apparatus with the head directed toward one of the closed arms. Mice were allowed to freely explore the apparatus for 10 min. The distance traveled, the number of the entries into each arm, and the time spent in each arm were automatically calculated with ANY-maze tracking software. In this behavioral experiment, the following exclusion criteria was adopted independently of the genotype or treatment (before the blind code was broken): mice that entered directly one of the closed arms from the starting central platform and spent the entire 10 minutes of the test in this closed arm were excluded from the analysis (4 out of 110 mice).

In vivo viral injection and micro optic implantation. All procedures were performed similarly to published protocols ^31, 93–95^. For stereotaxic surgery, animals were anesthetized with 2% isoflurane (Iso-Vet; Piramal Critical Care) and positioned into a digital stereotaxic frame (Stoelting) over a heating pad (∼37°C). After surgeries, the animals were maintained under a heating lamp until recovery. To reduce pain, inflammation and avoid infections, animals were administered with ketorolac (5 mg/Kg), dexamethasone (5 mg/Kg) and enrofloxacin (5 mg/Kg). During the first surgery, the viral vector expressing GCamP6f 96 was injected into the dorsal CA1 hippocampal region. In brief, after exposing the skull, a small hole was drilled at the following coordinates relative to bregma 97: antero-posterior: -1.94 mm; lateral: +1.20 mm. Animals were then injected with the GCaMP6f-expressing vector at -1.3 mm from the cortical surface using pulled glass capillaries connected to a Hamilton syringe mounted on a digital infusion micro-pump (Harvard Apparatus). Mice received 0.8 μl (flow rate: 0.1 µl/min) of Ready-to-Image virus AAV1.Camk2a.GCaMP6f.WPRE.bGHpA (Inscopix). After completing the infusion, the capillary was maintained in place for 10 minutes, and the skin was sutured upon the capillary removal. Upon recovery of the animals (3-4 weeks), a ProView™ Lens Probe (Inscopix) micro optic (1.0 mm diameter, 4.0 mm length) was implanted over the dorsal CA1 for imaging. To this aim, a 2.8 mm hole was drilled into the skull with a trephine drill over the stereotaxic coordinates where the GCaMP6f-expressing AAV was previously injected. To avoid compression of the CA1 by the lens, a cylindrical column of cortical tissue (∼2 mm diameter) and the upper part of the corpus callosum above the CA1 were aspirated with a 30G blunt-end needle, taking care of not damaging the tissue below. During surgery, the exposed brain tissue was constantly rinsed with ice-cold sterile PBS. Next, the nVista 3.0 miniaturized microscope (Inscopix) was used to place the lens probe over a suitable imaging field by assessing the overall fluorescence and the presence of blood vessels to be used as landmarks. Once completing the positioning, the exposed brain tissue around the base of the lens probe and the craniotomy were protected with Kwik-sil silicone (WPI). Then, the lens probe was permanently fixed to the skull with super-bond dental cement (Sun Medical). After an additional 4 weeks of animal recovery, a microscope magnetic baseplate was permanently fixed to the animal skull with the same dental cement. For this procedure, animals were lightly sedated with isoflurane and the miniaturized microscope was used again to guide the positioning of the baseplate over the lens probe to visualize the GCaMP6f-expressing neurons in the field of view. The magnetic baseplate attached to the skull allowed for repeated attachment and detachment of the microscope and thus the longitudinal imaging from the same CA1 field of view over multiple sessions. Between each imaging sessions, the microscope was removed and the top of the lens probe was protected with a magnetic cover sticking to the baseplate.

In vivo imaging sessions in the open field arena. Five days before starting the in vivo imaging experiments, mice where handled once a day for 5 min each day. Before the imaging experiments, mice were then habituated for 2-3 weeks to the microscope, wire and arena. During the first week of familiarization, mice were first habituated for one day to an empty arena made of grey opaque acrylic (44x44 cm) for 30 minutes. On the next two days of habituation, a ‘‘dummy’’ plastic scope (Inscopix) mimicking the shape and weight of the actual microscope was attached to the baseplate and again the animals were left exploring the arena for 30 minutes. During the following two weeks, the animals were daily treated with i.p. injections of vehicle (2% DMSO in saline) and also subjected to four additional habituation session of 1 hour each with the real microscope in order to familiarize to the microscope’s wire needed for data transfer. All habituation and imaging sessions were performed with dim light illumination (15-20 Lux).

To assess neuronal activity in the same neurons upon diazepam administration following vehicle or bumetanide treatment, we setup a longitudinal experiment consisting in two separate imaging sessions two weeks apart. During both imaging sessions, an ‘‘interrupted’’ imaging regime was used ^31^ to avoid potential photobleaching and/or phototoxicity deriving from exposure to the blue light needed to excite GCaMP6f fluorescence. The ‘‘interrupted’’ imaging regime consisted of active imaging periods (i.e. with blue light illumination) of 5 minutes alternated with 5 minutes of no light illumination.

On the first imaging session, animals were injected with vehicle (2% DMSO in saline) and placed after 10 minutes in the arena for 30 minutes during which neuronal calcium activity was recorded for a total of 15 minutes. Next, animals were injected with diazepam (Tocris; 2 mg/Kg in saline) and again placed after 10 minutes in the arena for 30 minutes to record neuronal imaging data for further 15 minutes. Following this first imaging section, animals were daily treated with i.p. injections of bumetanide (0.2 mg/Kg in saline) for two weeks. During this period, animals were also subjected to four additional habituation session of 1 hour each with the real microscope in order to maintain the experimental routine and habit to the microscope and wire. On the second imaging session, animals were injected with bumetanide (0.2 mg/Kg in saline) and placed after 10 minutes in the arena for 30 minutes during which neuronal calcium activity was recorded for 15 minutes. Finally, animals were injected with diazepam (2 mg/Kg in saline) and placed after 10 minutes in the arena for 30 minutes to record neuronal imaging data for further 15 minutes.

Processing and analysis of in vivo calcium imaging videos. Videos of calcium activity acquired by the miniature microscopes were processed with Inscopix data processing software (IDPS) following the manufacture recommendations. First, videos of each imaging sessions were spatially down-sampled by a factor of two and spatially band-passed (High cutoff: 0.5; Low cutoff: 0.005). Videos were also corrected for motion artifacts using an image registration method98 and transformed to relative changes in fluorescence (F/F0). The value of F/F0 for each pixel in the movie was calculated as (F-F0)/F0, where F0 is the mean fluorescence signal of the pixel during the entire recording. Cells were identified by principal component analysis (PCA) and independent component analysis (ICA), as previously described99, and the corresponding calcium traces were extracted. Next, the calcium traces from the two imaging sessions (vehicle plus diazepam, and bumetanide plus diazepam) were longitudinally concatenated in time. Traces were manually inspected to remove cells showing more than one independent component. Finally, calcium event features (i.e. starting time and amplitude) were automatically detected using changes in F/F0 of at least 8 units of median absolute deviation (MAD) and a minimal decay time constant ( ) of 100 ms 96.

The calcium event data were then analyzed with custom scripts in MATLAB (Mathworks). Cells presenting a signal-to-noise-ratio lower than 6 were excluded from the analysis. This threshold allowed the removal of neurons with low quality in terms of signal-to-noise-ratio or background fluctuations. To determine the cut-off parameter for exclusion, the overall distribution of the SNR for all recorded neurons was plotted and the threshold values selected to remove values less than the 10th percentile of the distribution. To quantify the level of activity, the mean events rate (MER, events/sec), defined as the mean number of calcium events in the diverse recording time windows, was computed. Neurons presenting a MER lower than 0.01 events/sec were excluded from the analysis to eliminate silent cells or cells showing very low activity. The same parameters were used for all animals. As previously described for cultures and slices, the MER ratio calculated as the MER during diazepam treatment divided by the MER during vehicle or bumetanide treatment was calculated. To quantify the mixed population of neurons the bootstrap methods used for cultures was adapted to the calcium imaging data. To this aim, four null distributions reflecting the overall changes in MER for all neurons in the four different experimental conditions (i.e., WT and Ts65Dn mice pretreated with vehicle plus diazepam or bumetanide plus diazepam) were computed as it follows. For each neuron belonging to the four experimental conditions, the event trains for each time segment (vehicle plus diazepam and bumetanide plus diazepam) was divided into 1-min bins. The MERs recorded during each 1-min bin for the two time segments for each neuron were randomly shuffled into two groups for 10,000 times. The differences between the MER of the randomly shuffled groups belonging to the same experimental conditions were used to generate the four corresponding null distributions. Then, for each neuron, the real difference between the MER during vehicle or bumetanide and the MER during vehicle plus diazepam or bumetanide plus diazepam was computed, and assessed whether it fell outside the 85% confidence interval of the corresponding null distribution. Thus, the overall percentage of neurons showing significant changes in the MER and belonging to the three different categories (i.e., increase, decrease or not change) was computed for each experimental condition.

To identify the populations of neurons with different responses to bumetanide and/or diazepam treatment, two different ratios for each neuron were computed: i) ratio-1: the MER during bumetanide divided by the MER during vehicle; ii) ratio-2: the MER during diazepam plus bumetanide divided by the MER during diazepam plus vehicle. The first ratio reflected the variation in the neuronal activity elicited by bumetanide treatment in comparison to vehicle, while the second ratio returned the variation in the neuronal activity upon bumetanide plus diazepam administration in comparison to vehicle plus diazepam treatment. The two ratios were correlated in a scatterplot (i.e. ratio-1 on x-axis, ratio-2 on y-axis). The plot was divided into four quadrants based on the possible different neuronal activity in response to diazepam or bumetanide treatment over vehicle. Cells showing a decrease in the MER after both bumetanide and bumetanide plus diazepam treatments fell in quadrant 1 (decrease in both ratios). Cells showing an increase in MER after bumetanide and a decrease after bumetanide plus diazepam administrations fell in quadrant 2 (increase in ratio-1, decrease in ratio-2). Cells showing an increase in MER after both bumetanide and bumetanide plus diazepam administrations fell in quadrant 3 (increase in ratio-1, increase in ratio-2). Cells showing a decrease in MER after bumetanide and an increase after bumetanide plus diazepam administration fell in quadrant 4 (decrease in ratio-1, increase in ratio-2). Cells showing MER ratios changes below 10% were considered non-responsive to treatments and were excluded from further analysis. To predict the effect of diazepam plus bumetanide elicited in each Ts65Dn neuron based on MER variations during the previous sessions, a machine learning approach was implemented. For each neuron, six different features including MER, mean, minimum and maximum amplitude of the Ca2+ transients, mean and maximum distance between Ca2+ events were calculated. These features were computed in the first three temporal concatenated experimental phases (i.e., vehicle, diazepam and bumetanide) while the features in bumetanide plus diazepam phase were hidden to the classification algorithm. Then, the Ts65Dn dataset was randomly divided in the training and test sets, representing respectively the 54% and 46% of the total dataset. For simplicity, the machine learning classificatory was asked to predict only two possible output labels (categories: an increase (>1) or a decrease (<1) of the MER during diazepam plus bumetanide treatment, when compared to the diazepam plus vehicle. Both datasets contained a comparable percentage of neurons belonging in the two categories (training set: 65% of <1 category and 35% of >1 category; test set: 58% <1 category and 45% >1 category). Supporting Vector Machine classificator (SVM, 100) was trained to predict the categorical label (<1 or >1) of the neuron from the feature vectors. The accuracy of the classificator in the test set was tested using the Matlab function “classperf”, which computed the following parameters: the correct rate (i.e., number of correctly classified samples divided by the number of classified samples), the error rate (i.e., number of incorrectly classified samples divided by the number of classified samples), sensitivity (i.e., number of correctly classified positive samples divided by the number of true positive samples), specificity (i.e., number of correctly classified negative samples divided by the number of true negative samples), positive predictive value (i.e., number of correctly classified positive samples divided by the number of positive classified samples), negative predictive value (i.e., number of correctly classified negative samples divided by the number of negative classified samples), predicted likelihood (1 - Sensitivity/Specificity) and prevalence (number of true positive samples divided by the total number of samples; Supplementary Table 2).

Statistical analysis. Except were otherwise stated, the results are presented as the means ± SEM. Statistical analysis was performed using SigmaPlot (Systat) or GraphPad (Prism) software. Where appropriate, the statistical significance was assessed using the following parametric test: Student’s t test or two-way ANOVA followed by all pairwise Turkey’s post hoc test. In case normal distribution or equal variance assumptions were not valid, statistical significance was evaluated using the following non parametric tests: Mann-Whitney Rank Sum Test or two-way ANOVA on ranked transformed data. For statistical evaluation of in vivo calcium imaging longitudinal data, we used the linear mixed model (LME) from nlme package in R (http://CRAN.R-project.org), as recently suggested101. Specifically, we fitted the LME using genotype and treatment as fixed effects, and cell number and mouse ID as nested random effects. Statistical difference in the percentage of neurons (Fig. 4 and 5) were analyzed by Chi-squared test with Sidak adjustment for multiple comparisons. For all tests, P values < 0.05 were considered significant.

## Acknowledgements

The authors wish to thank Valentina Pasquale, IIT, Italy for her help with the MEA data analysis and Giacomo Pruzzo, IIT, Italy for constructing the chamber for long-term MEA measurement and the apparatus used for *in vivo* calcium imaging in freely moving animals. We thank the IIT animal facility staff for their valuable work. We also thank the members of the RNA initiative at IIT for the helpful discussions. This work was partially funded by the European Research Council (ERC) under the European Union’s Horizon 2020 research and innovation program (Grant Agreement No. 725563 to L.C.), Telethon Foundation (TCP15021 to L.C.), the Jerome Lejeune Foundation (EPFD0149 to IC) and Angelini Foundation (168(A)MD21320 to LC).

## Author Contributions

I.C. carried out the electrophysiological experiments with microelectrode arrays, analysed the data, prepared the figures and wrote the manuscript. Surgeries and *in vivo* calcium imaging experiments in freely moving animals were performed by A.C. while the I.C. analysed the related data. M.R. prepared human iPSC-derived neurons and collected and analysed the biochemical and calcium imaging data of iPSC-derived neurons. M.P. collected and analysed the behavioural data. M.A. and A.C. collected and analysed the chloride imaging and the biochemical data. M.A. and M.N. prepared primary neuronal cultures. A.C. performed and analysed the calcium imaging experiments in primary neuronal culture. M.C. supervised the experiments with the microelectrode array. A.C. conceived the study, designed and supervised the experiments, and wrote the manuscript. L.C. conceived the study, supervised the experiments and wrote the manuscript. All authors read and revised the manuscript.

## Conflicts of Interest

LC is the cofounder and a scientific advisor at IAMA Therapeutics. LC and AC are inventors on the following patents: US 9,822,368 (granted 2017); EP 3083959 (granted 2019); JP 6490077 (granted 2019) and US 11427836 (granted 2022), US 17/861676, EP 18717045.1, HK 62020014163.3, CA 3059389, CN 201880038547.7, JP 2020-505522, and IL 269952.

