## Supplementary Figure for "Heterogeneous subpopulations of GABAAR-responding neurons coexist in physiological and pathological mature neuronal networks at increasing scales of complexity"

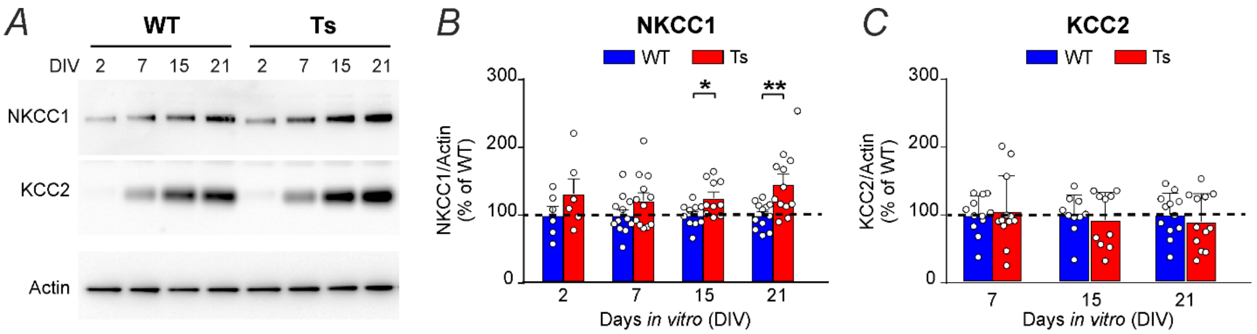


**Supplementary Figure 1. NKCC1 protein level is increased in Ts65Dn neurons in culture during *in vitro* development. A)** Representative immunoblots for NKCC1 and KCC2 protein extracts from WT and Ts65Dn cortical neuronal cultures at different stages of development. Actin was used as an internal standard. Full blots are shown in Supplementary Fig. 10B. **B)** Quantification of average NKCC1 protein (± SEM) expressed as the percentage of WT neurons (dotted line) at each time point in experiments as in A. **C)** Quantification of average KCC2 protein (± SEM) in the same experiments in B. KCC2 at 2 DIV was not quantified because expression was almost undetectable. Dots indicate values of individual culture wells (obtained from 5 independent neuronal cultures). *p<0.05, **p<0.01, Student's t test or Mann-Whitney Rank Sum Test.

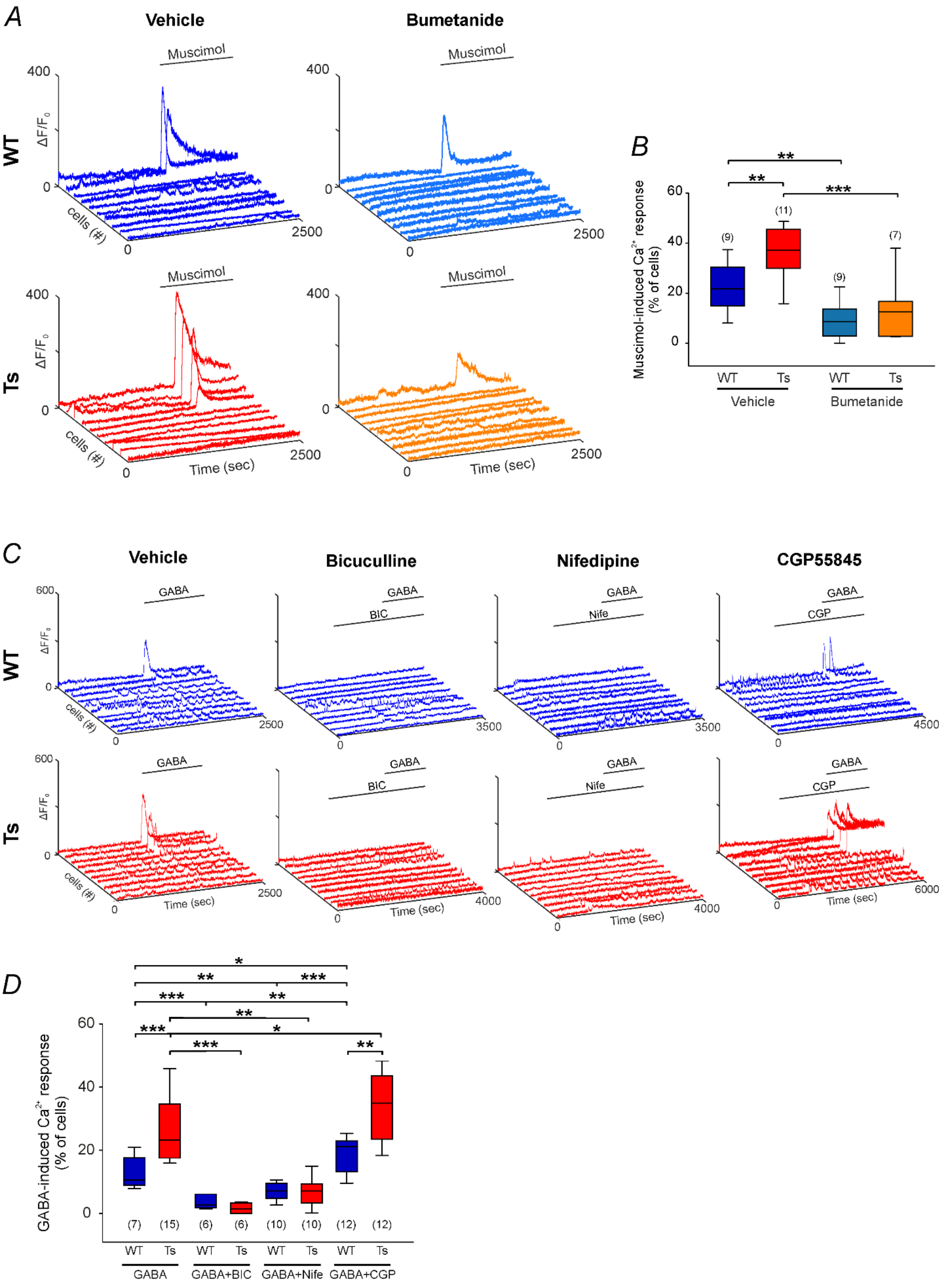
A
Vehicle
Bumetanide
Muscimol
WT
Ts
ΔF/F0
400
0
cells (#)
2500
Time (sec)
B
Muscimol-induced Ca2+ response (% of cells)
60
40
20
0
(9)
(11)
(7)
**
***
WT
Ts
Vehicle
Bumetanide
C
Bicuculline
Nifedipine
CGP55845
GABA
BIC
Nife
CGP
600
3500
4000
4500
6000
D
GABA-induced Ca2+ response (% of cells)
*
(15)
(6)
(10)
(12)
GABA+BIC
GABA+Nife
GABA+CGP

**Supplementary Figure 2. GABA-induced depolarizing responses are mediated by $GABA_A$Rs and L-type voltage-gated $Ca^{2+}$ channels. A)** Representative calcium traces of hippocampal neuronal cultures at 15 DIV, loaded with the $Ca^{2+}$-sensitive dye Fluo-4 upon bath application of the specific $GABA_A$R agonist muscimol (10 μM). Neurons were pretreated with vehicle (0.01% DMSO) or the NKCC1 inhibitor bumetanide (10 μM). **B)** Quantification of the percentage of neurons showing depolarizing muscimol-induced responses in experiments as in A. Boxplots indicate median and 25th-75th percentiles, whiskers represent the 5th-95th percentiles. Numbers in parenthesis indicate the number of analyzed coverslips for each experimental group (obtained from 3 independent neuronal cultures). **$P<0.01$, ***$P<0.001$, Tukey *post hoc* test following two-way ANOVA. **C)** Representative calcium traces of hippocampal neurons at 15 DIV, loaded with the $Ca^{2+}$-sensitive dye Fluo-4 upon bath application of GABA (100 μM). Cells at 15 DIV were pretreated with either vehicle (0.01% DMSO), the $GABA_A$ receptor antagonist bicuculline (BIC; 100 μM), the L-type voltage-gated $Ca^{2+}$ channels blocker nifedipine (Nife; 10 μM) or the $GABA_B$ receptor antagonist CGP55845 (CGP; 10 μM). **D)** Quantification of the percentage of neurons showing depolarizing GABA responses in experiments as in C. Boxplots indicate the median and 25th-75th percentiles, whiskers represent the 5th-95th percentiles. Numbers in parenthesis indicate the number of analyzed coverslips for each experimental group (obtained from 2-6 independent neuronal cultures). *$p<0.05$, **$p<0.01$, ***$p<0.001$, Tukey *post hoc* test following two-way ANOVA.

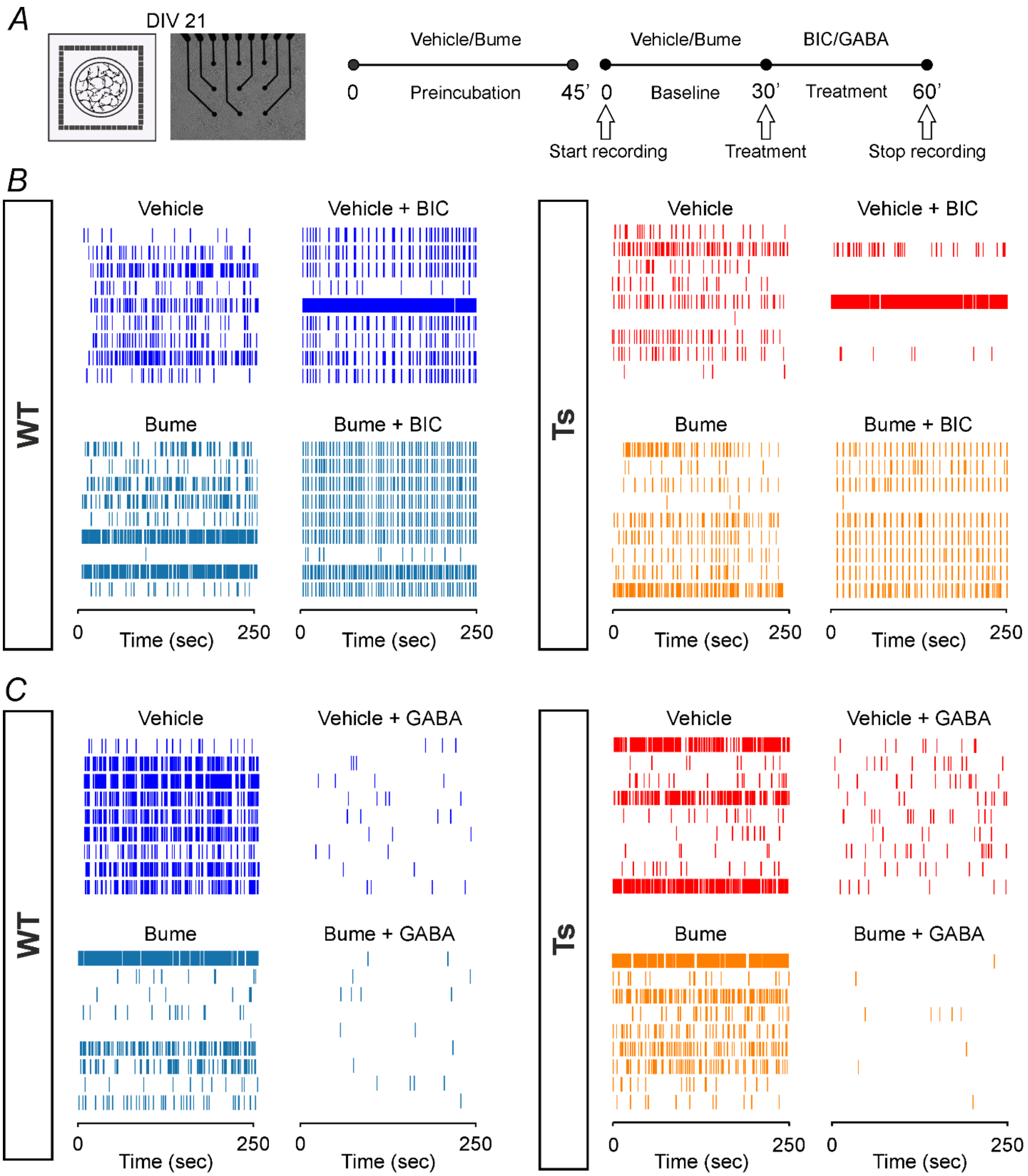
A
DIV 21
Vehicle/Bume
Vehicle/Bume
BIC/GABA
0
Preincubation
45'
0
Baseline
30'
Treatment
60'
Start recording
Treatment
Stop recording
B
WT
Vehicle
Vehicle + BIC
Bume
Bume + BIC
Ts
Vehicle
Vehicle + BIC
Bume
Bume + BIC
0
Time (sec)
250
C
WT
Vehicle
Vehicle + GABA
Bume
Bume + GABA
Ts
Vehicle
Vehicle + GABA
Bume
Bume + GABA
0
Time (sec)
250

**Supplementary Figure 3**. **Mixed subpopulations of neurons with hyperpolarizing or depolarizing $GABA_AR$ signaling are present in WT and Ts65Dn mature neuronal cultures grown over MEA. A)** *Left*: Schematic representation of primary hippocampal neuronal cultures grown over a microelectrode array (MEA) for electrophysiological recordings. *Center*: Representative transmitted-light image showing an MEA seeded with hippocampal neurons at 21 DIV. *Right*: Schematic representation of the experimental protocol; neuronal cultures were pre-incubated with vehicle (0.01% DMSO) or bumetanide (10 μM) for 45 minutes followed by 30 minute recording of spontaneous activity. Neurons were then recorded for additional 30 minutes, after addition of bicuculline (BIC; 20 μM) or GABA (100 μM). **B, C)** Examples of 250-second raster plots of spiking activity of WT and Ts65Dn neuronal cultures after bath application of the indicated drugs. Data are examples derived from the experiments shown as averages in Figure 2.

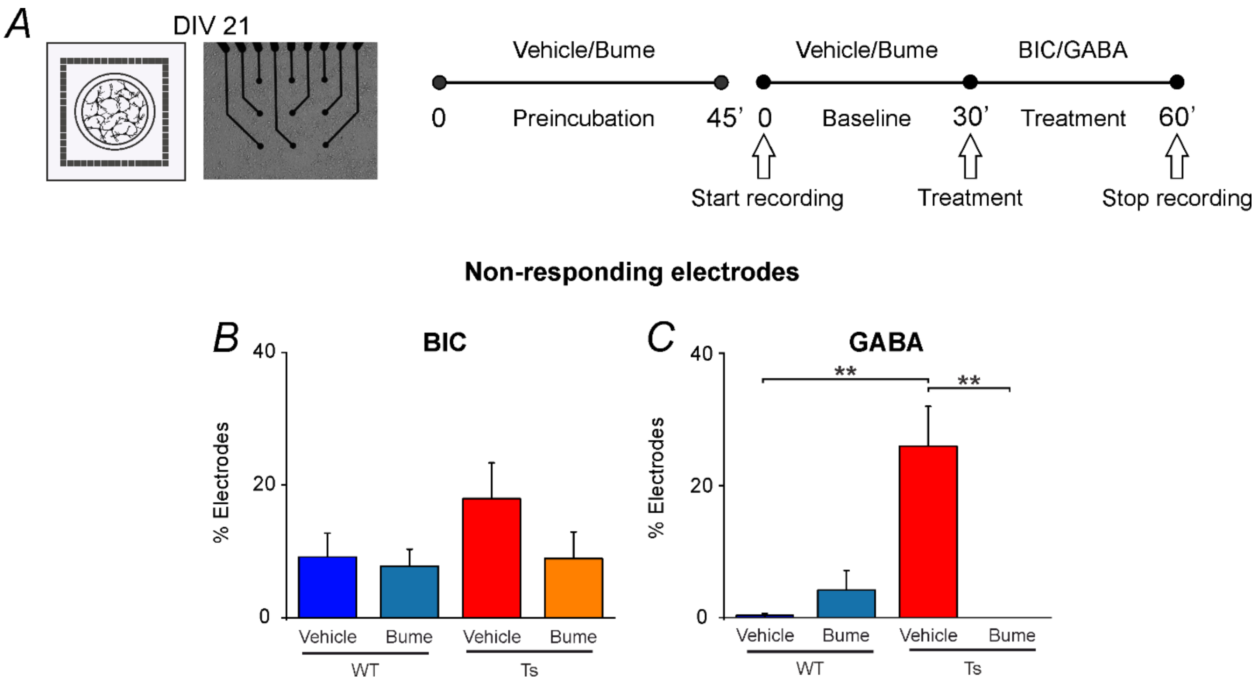


**Supplementary Figure 4**. **Ts65Dn hippocampal mature cultures show an increased number of GABA non-responding electrodes compared to WT cultures, and this number decreases upon bumetanide administration. A)** *Left*: Schematic representation a of primary hippocampal neuronal culture grown over a microelectrode array (MEA) for electrophysiological recordings. *Center*: Representative transmitted-light image showing a MEA seeded with hippocampal neurons at 21 DIV. *Right*: Schematic representation of the experimental protocol; neuronal cultures were pre-incubated with vehicle (0.01% DMSO) or bumetanide (10 µM) for 45 minutes followed by 30 minute recording of spontaneous activity. Neurons where then recorded for additional 30 minutes, after addition of bicuculline (BIC; 20 µM) or GABA (100 µM). **B**, **C**) Percentage of electrodes that did not change significantly the MFR upon BIC (B) or GABA (C) treatment for WT and Ts65Dn mature cultures in the experiments shown in Figure 2. ** $p<0.01$; Tukey's *post hoc* test following two-way ANOVA.

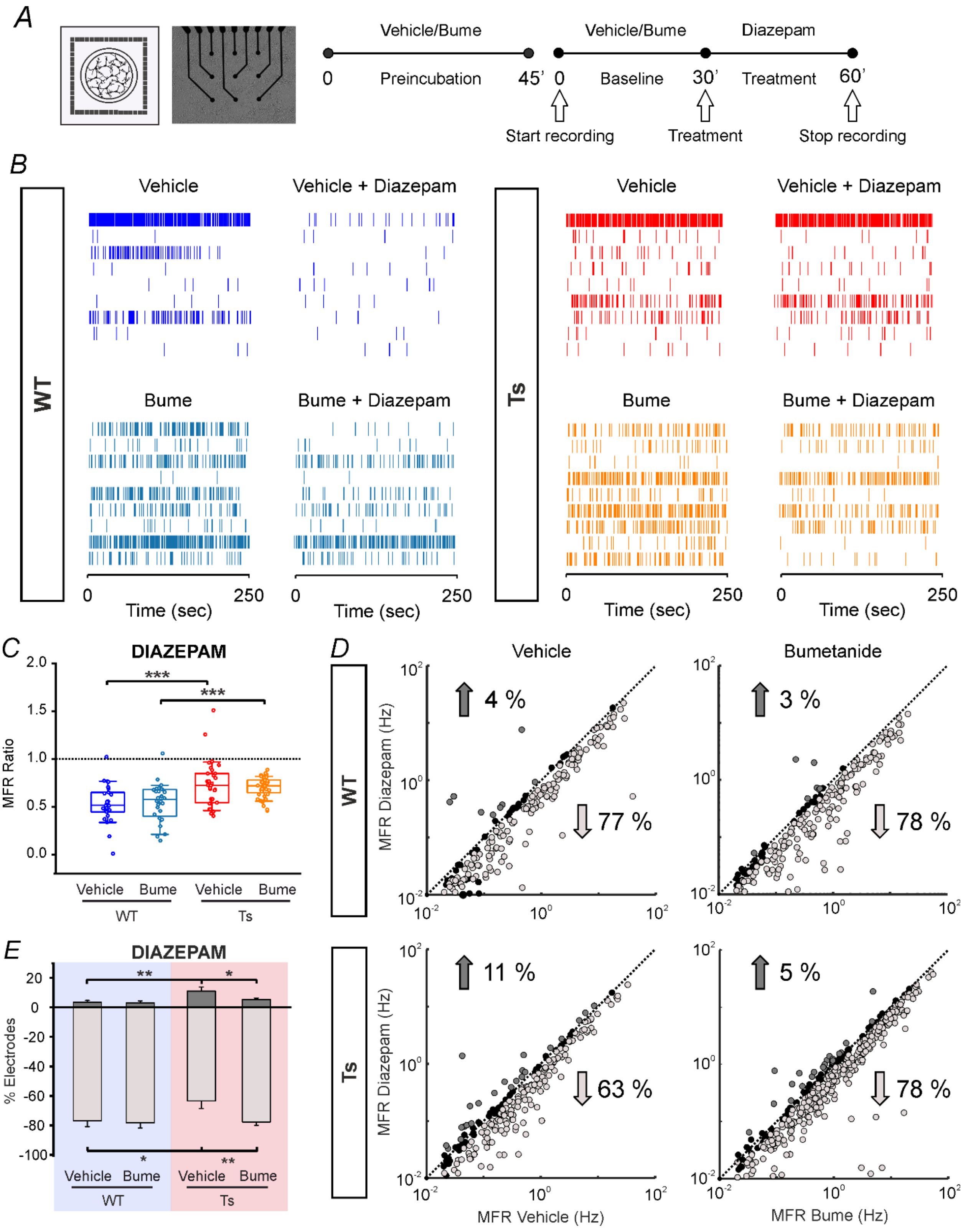
A
Vehicle/Bume
Preincubation
0
45'
Vehicle/Bume
Diazepam
0
Baseline
30'
Treatment
60'
Start recording
Treatment
Stop recording
B
WT
Vehicle
Vehicle + Diazepam
Bume
Bume + Diazepam
Ts
Time (sec)
0
250
C
DIAZEPAM
MFR Ratio
2.0
1.5
1.0
0.5
0.0
***
***
Vehicle
Bume
WT
Ts
D
Vehicle
Bumetanide
MFR Diazepam (Hz)
4 %
77 %
3 %
78 %
11 %
63 %
5 %
78 %
MFR Vehicle (Hz)
MFR Bume (Hz)
E
DIAZEPAM
% Electrodes
20
0
-20
-40
-60
-80
-100
**
*
**

**Supplementary Figure 5**. **Bumetanide treatment rescues the paradoxical effect of diazepam on Ts65Dn neuronal networks cultured over MEAs. A)** *Left*: Schematic representation of primary hippocampal neurons in cultures grown over a microelectrode array (MEA) for electrophysiological recordings. *Center*: Representative transmitted-light image showing an MEA seeded with hippocampal neurons at 21 DIV. *Right*: Schematic representation of the experimental protocol; neuronal cultures were pre-incubated with vehicle (0.01% DMSO) or bumetanide (10 µM) for 45 minutes followed by 30 minute recording of spontaneous activity. Neurons where then recorded for additional 30 minutes after addition of diazepam (1 µM). **A, B)** Examples of 250-second raster plots of spiking activity of WT and Ts65Dn neuronal cultures after bath application of the indicated drugs. **C)** Quantification of the mean firing rate (MFR) ratio of WT and Ts65Dn neuronal cultures in same experiments as in B. MFR ratio over baseline firing (dotted line) was calculated for each electrode and then averaged for each MEA. In the boxplot, the small square indicates the mean, the central line illustrates the median, the box limits indicate the 25th and 75th percentiles, the whiskers represent the 5th-95th percentiles and each dot represents the MFR ratio for each recorded culture. Numbers in parenthesis indicate the number of analyzed MEA for each experimental group (obtained from 8 independent neuronal cultures). *** $p<0.001$; Tukey's *post hoc* test following two-way ANOVA. **D**) Scatter plots showing the MFR for each active electrode (plotted as a dot) from all recorded MEAs (shown in C as ratio) before (x-axis) and after (y-axis) bath application of diazepam. Dark-grey dots represent electrodes showing a significant increase in MFR. Light-grey dots represent electrodes showing a significant decrease in the MFR. Black dots represent electrodes showing no significant changes in MFR. Significant changes in MFR (numbers with arrows) for each electrode upon diazepam application were evaluated by bootstrap analysis. **E)** Quantification of the average percentage number (± SEM) of MEA electrodes (in the same experiments showed in C), displaying significant changes in MFR by bootstrap analysis after diazepam administration in comparison to their basal condition in WT (blue) and Ts65Dn (pink) neuronal cultures. * $p<0.05$; ** $p<0.01$; Tukey's *post hoc* test following two-way ANOVA.

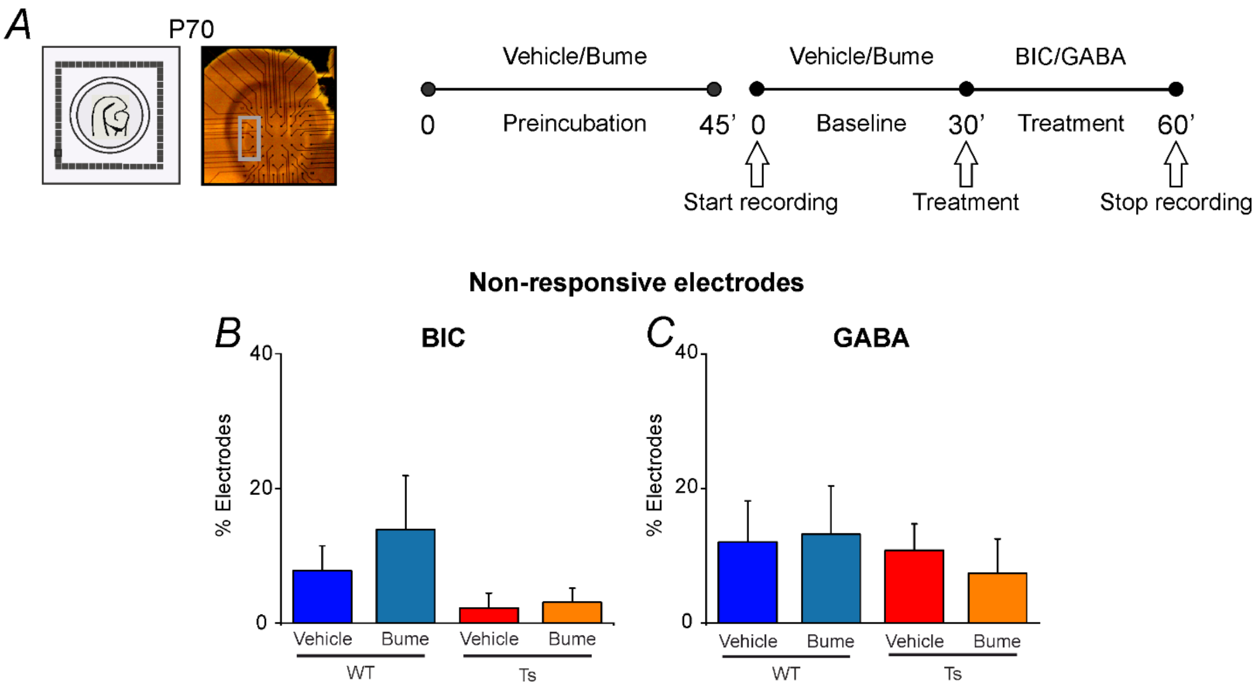


**Supplementary Figure 6**. **The percentage of GABA or BIC non-responding electrodes did not change between brain slices from adult WT and Ts65Dn mice. A)** *Left*: Schematic representation of an acute brain slice of the hippocampus-entorhinal cortex (EC) region from an adult animal for electrophysiological recordings. The slice is positioned on an MEA for recording. *Center*: Representative picture of the experimental setup with MEA electrodes and overlying hippocampus-EC slice. The grey square highlights the analyzed electrodes positioned in the CA1 region. *Right*: Schematic representation of the experimental protocol. Brain slices were pre-incubated with vehicle (0.01% DMSO) or bumetanide (10 µM) for 45 minutes followed by 30 minute recording of spontaneous activity. Brain slices where then recorded for additional 30 minutes, after addition of bicuculline (BIC; 20 µM) or GABA (100 µM). **B, C)** Percentage of electrodes that did not change significantly the MFR upon BIC (B), or GABA (C) treatment in hippocampal slices from adult WT and Ts65Dn mice in the same experiments shown in Figure 3.

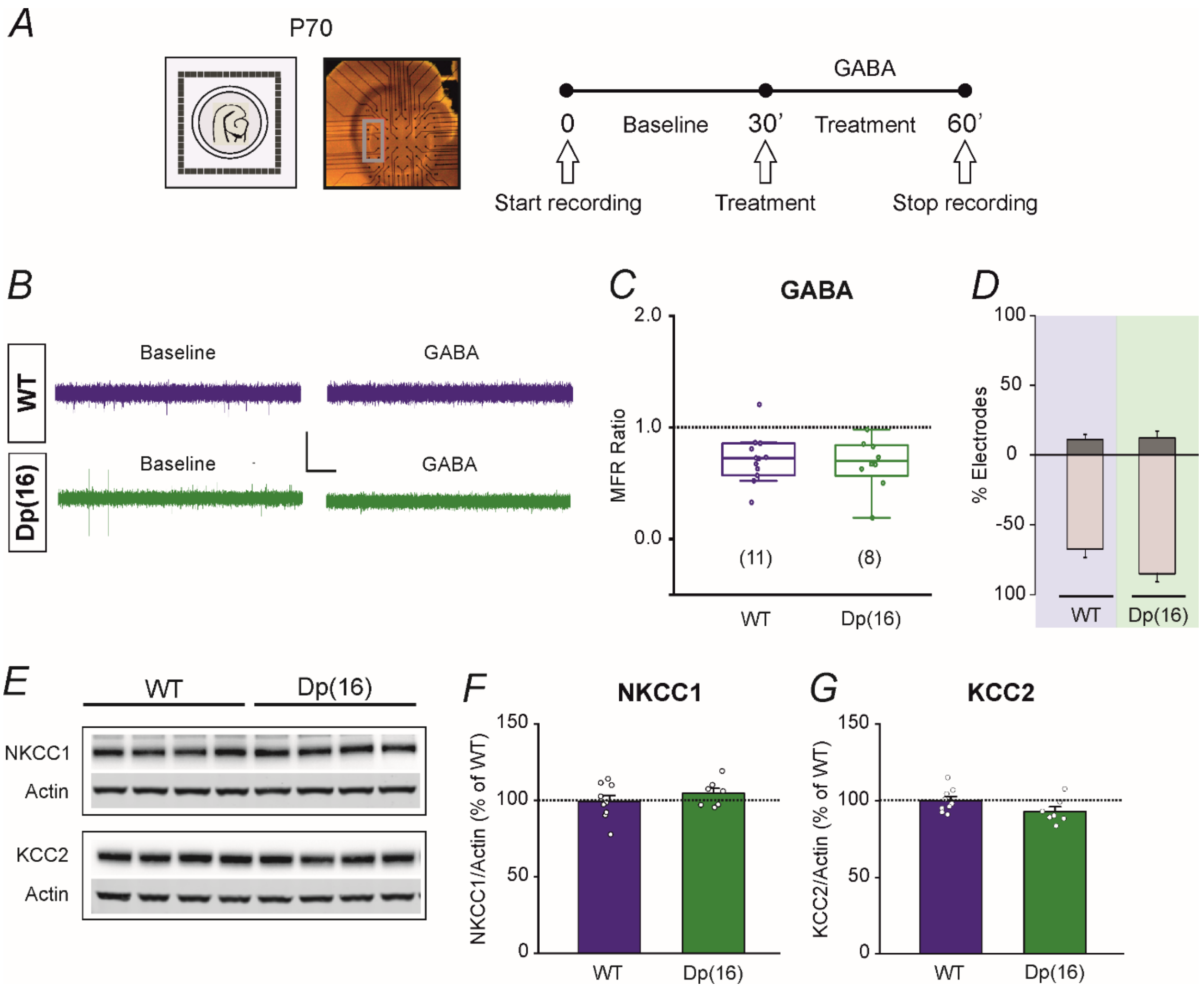
A
P70
GABA
0
Baseline
30'
Treatment
60'
Start recording
Treatment
Stop recording
B
WT
Dp(16)
Baseline
GABA
Baseline
GABA
C
GABA
2.0
1.0
0.0
MFR Ratio
(11)
(8)
WT
Dp(16)
D
100
50
0
-50
100
% Electrodes
WT
Dp(16)
E
WT
Dp(16)
NKCC1
Actin
KCC2
Actin
F
NKCC1
150
100
50
0
NKCC1/Actin (% of WT)
WT
Dp(16)
G
KCC2
150
100
50
0
KCC2/Actin (% of WT)
WT
Dp(16)

**Supplementary Figure 7. Adult WT (C57BL/6J) mice and the Dp(16)1Yey/+) mouse model of DS confirmed mixed subpopulations of GABA hyperpolarizing or depolarizing neurons, whereas Dp(16)1Yey/+ DS mice did not show an increased population of depolarizing GABA neurons or altered expression of chloride transporters. A)** *Left*: Schematic representation of an acute brain slice of the hippocampus-entorhinal cortex region from a P70 adult animal. The slice is positioned on an MEA for electrophysiological recordings. *Center*: Representative picture of the experimental setup with MEA electrodes and an overlying hippocampus-EC slice. The gray square highlights the analyzed electrodes, which were positioned in the CA1 region. *Right*: Schematic representation of the experimental protocol. We recorded 30 minutes of spontaneous activity (baseline) followed by 30 minutes of recording upon bath application of GABA (100 μM). **B)** Representative high-pass filtered (>300 Hz) recording traces for slices obtained from adult (3-4 months) WT (C57BL/6J) mice and the Dp(16)1Yey/+ mouse model of DS, and recorded before (baseline) and after bath application of GABA. Scale bars: 50 μV, 1 s; **C)** Quantification of the MFR ratio of WT and Dp(16)1Yey/+ slices upon GABA bath application. MFR ratio over baseline firing (dotted line) was calculated for each electrode and then averaged for each MEA. In the boxplot, the small square indicates the mean, the central line illustrates the median, the box limits indicate the 25th and 75th percentiles, the whiskers represent the 5th-95th percentiles and each dot represents the MFR ratio for each recorded slice. Numbers in parenthesis indicate the number of analyzed slices for each experimental group. **D)** Quantification of the average percentage number (± SEM) of MEA channels (in the same experiments in C), showing changes (by at least 15%) in the MFR after GABA administration in WT (blue) and Dp(16)1Yey/+ (green) slices. **E)** Representative immunoblots for NKCC1 and KCC2 protein extracts from hippocampi of adult WT and Dp(16)1Yey/+ animals. Actin was used as an internal standard. **F)** Quantification of NKCC1 protein in WT and Dp(16)1Yey/+ hippocampi expressed as the percentage of WT in experiments as in E. **G)** Quantification of KCC2 protein in WT and Dp(16)1Yey/+ hippocampi expressed as the percentage of WT in the same experiments shown in F. In F and G, data are expressed as means (±SEM). Dots indicate values of individual samples. Actin was used as an internal standard. Full blots are shown in Supplementary Fig. 10 C.

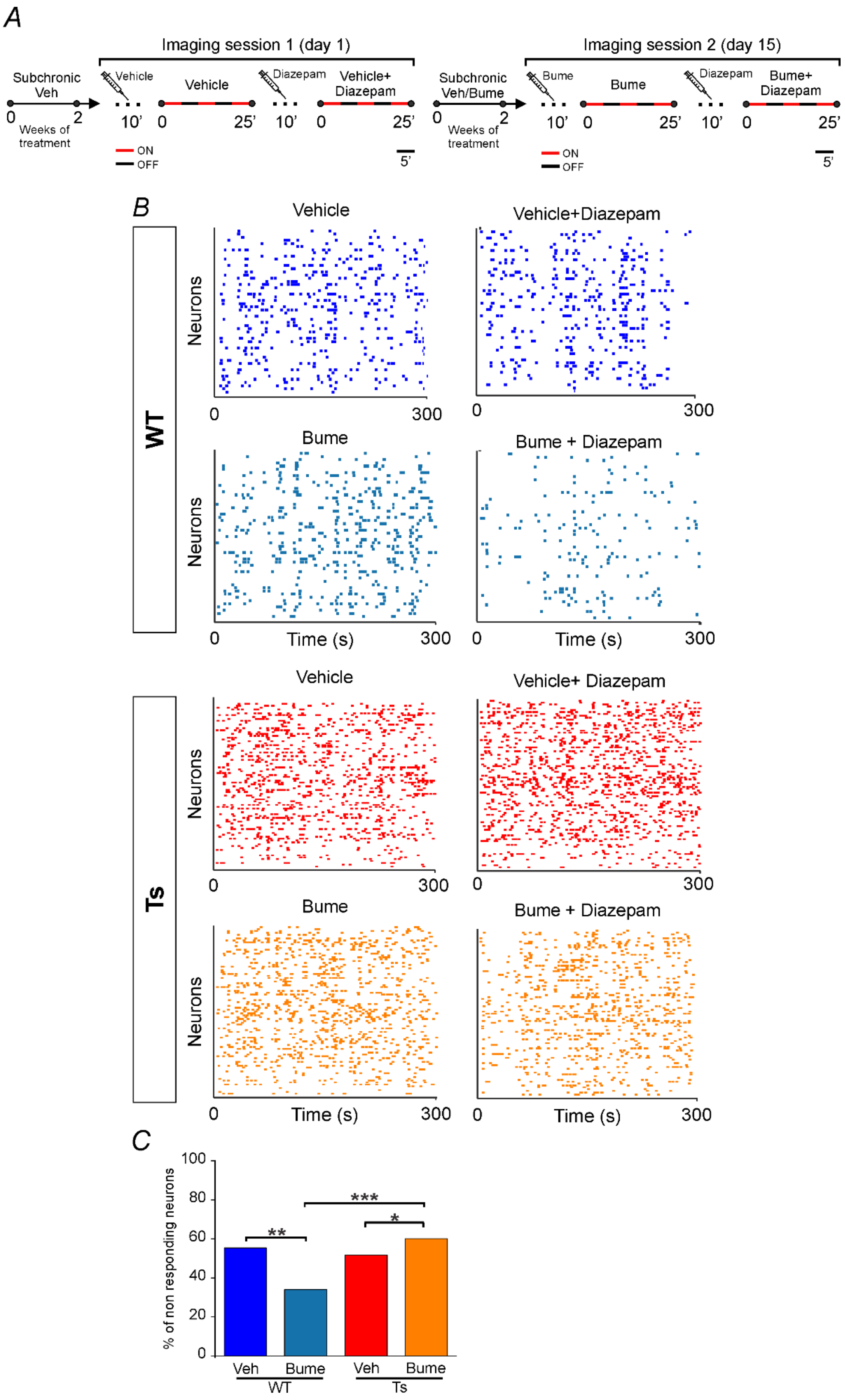

A
Imaging session 1 (day 1)
Subchronic Veh
Vehicle
Vehicle
Diazepam
Vehicle+ Diazepam
0
2
Weeks of treatment
10'
0
25'
10'
0
25'
ON
OFF
5'
Imaging session 2 (day 15)
Subchronic Veh/Bume
Bume
Bume
Diazepam
Bume+ Diazepam
0
2
Weeks of treatment
10'
0
25'
10'
0
25'
ON
OFF
5'
B
WT
Vehicle
Vehicle+Diazepam
Bume
Bume + Diazepam
Neurons
Time (s)
0
300
Ts
Vehicle
Vehicle+ Diazepam
Bume
Bume + Diazepam
Neurons
Time (s)
0
300
C
% of non responding neurons
100
80
60
40
20
0
**
***
*
Veh
Bume
Veh
Bume
WT
Ts

**Figure Supplementary 8**. **Bumetanide treatment partially rescues the depolarizing $GABA_AR$-mediated signaling in adult Ts65Dn mice, and differentially affects the number of diazepam non-responding-neurons in WT *vs* Ts65Dn mice *in vivo*. A)** Schematic representation of the experimental protocol timeline in WT and Ts65 adult (2-3 month old) mice *in vivo*. $Ca^{2+}$ events were imaged in the same neuron in two consecutive sessions before and after administration of the $GABA_AR$ positive allosteric modulator diazepam (2 mg/Kg), following a sub-chronic (2 weeks) treatment with either vehicle (2% DMSO in saline) or bumetanide (0.2 mg/Kg). Within each imaging session, neuronal activity was recorded in periods of 5 minutes ("ON", red) alternated to 5 minutes of rest ("OFF", blue) to collect 15 min of data for each session, while avoiding phototoxicity. **B)** Example of 5 minute raster plots of $Ca^{2+}$ events of CA1 neurons from WT and Ts65Dn mice recorded *in vivo* after diazepam treatment in the same experiments shown in Figure 4. **C)** Percentage of CA1 neurons that did not significantly change the MER upon diazepam treatment in adult WT and Ts65Dn mice in the same experiments shown in Figure 4. * $p<0.05$, ** $p<0.01$, *** $p<0.001$, Chi-Square test with Sidak adjustment for multiple comparisons.

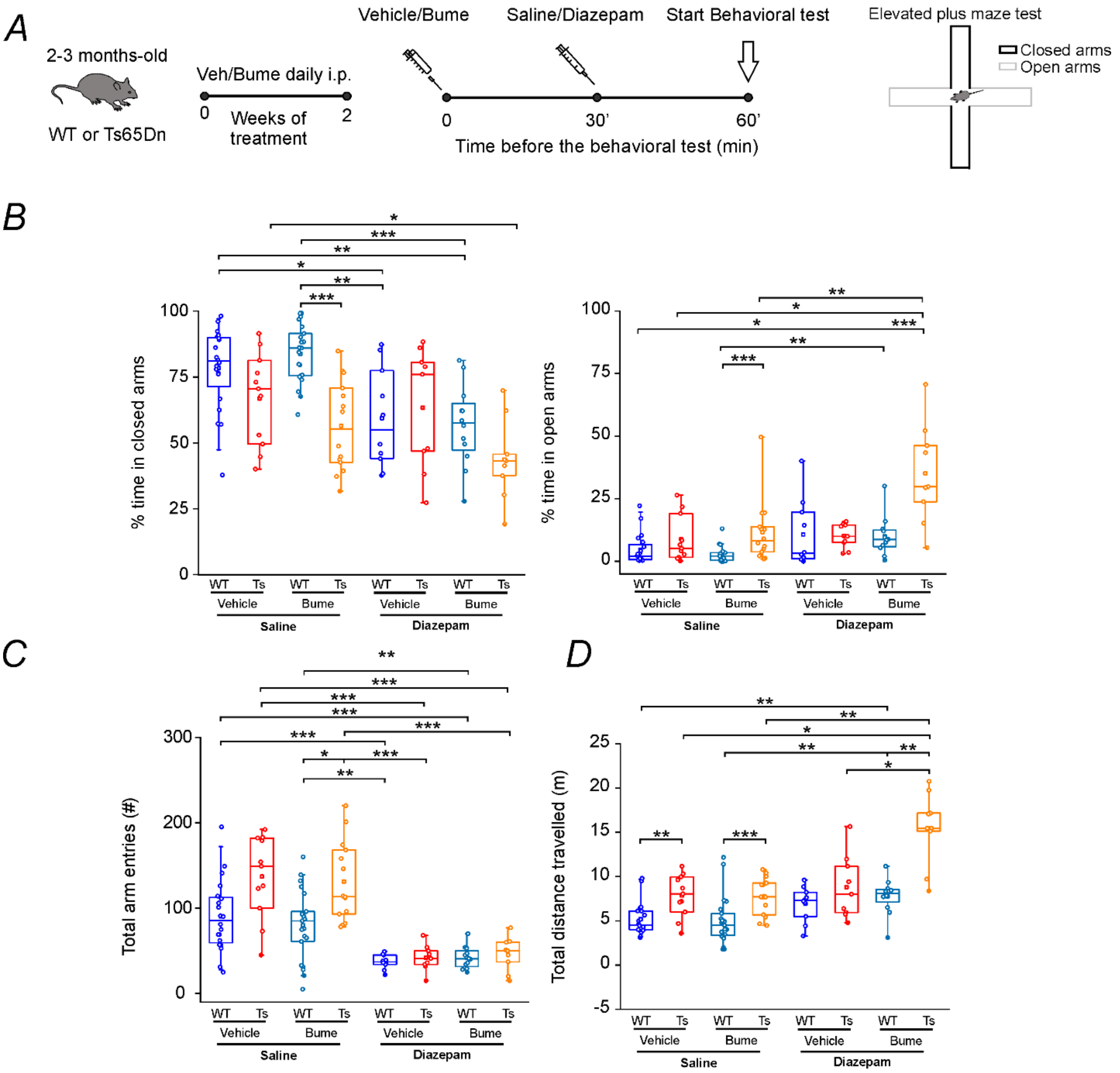


**Figure Supplementary 9. Diazepam treatment does not induce paradoxical anxiety responses, but bumetanide treatment potentiates diazepam anxiolytic action in Ts65Dn adult mice performing the elevated plus maze test. A)** Schematic representation of the experimental protocol. Adult WT and Ts65Dn mice were treated with bumetanide (0.2 mg/Kg i.p.) or the corresponding vehicle (2% DMSO in saline) for 2 weeks and then assessed in the elevated plus maze test. On the day of the test, mice were pretreated with vehicle or bumetanide and then tested 30 minutes after diazepam (2 mg/Kg) or saline administration. **B)** Quantification of the percentage time spent in the open (*Left*) and closed (*Right*) arms of the elevated plus maze for WT and Ts65Dn mice. **C)** Quantification of the total number of arm entries in the elevated plus maze for WT and Ts65Dn mice. **D)** Quantification of the total distance traveled in the elevated plus maze for WT and Ts65Dn mice. In all boxplots, the small square indicates the mean, the central line illustrates the median, the box limits indicate the 25th and 75th percentiles, the whiskers represent the 5th-95th percentiles and each dot indicates a value obtained from an individual animal. *p<0.05, ** p<0.01, *** p<0.001; Tukey's *post hoc* test, following two-way ANOVA.

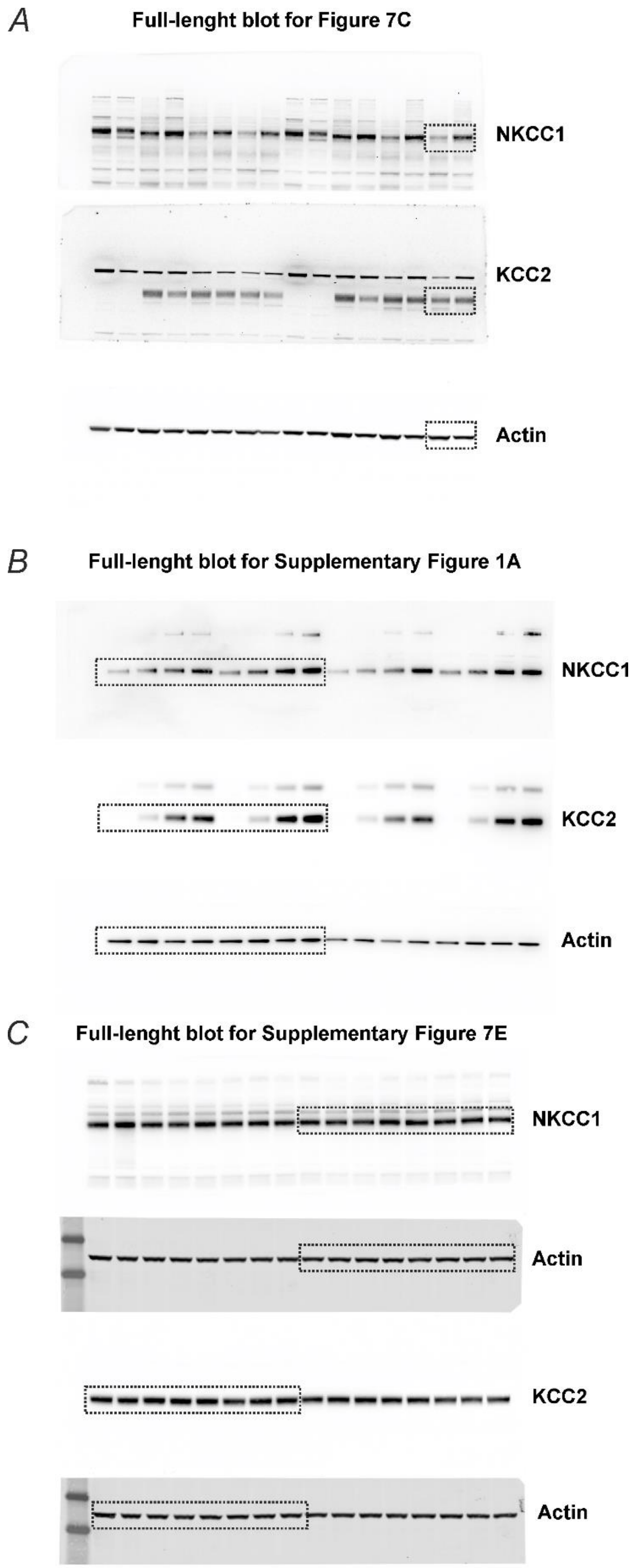


**Figure Supplementary 10.** Full-length blot images with dotted squared areas corresponding to the cropped western blot presented in Figure 7C and Supplementary Figure 1A and 7E.

| Parameters | Value |
|---|---|
| Correct Rate | 0.6378 |
| Error Rate | 0.3622 |
| Sensitivity | 0.6104 |
| Specificity | 0.6574 |
| Positive Predictive Value | 0.5595 |
| Negative Predictive Value | 0.7030 |
| Positive Likelihood | 1.7817 |
| Negative Likelihood | 0.5926 |
| Prevalence | 0.4162 |

**Supplementary Table 1. Performance-related parameters of the Supporting Vector Machine (SVM) classifier used in Figure 5.**

| Model | Reference | Structure | DIV/Age | GABAergic agonist/antagonist(concentration) | Technique | Reported neurons with depolarizing GABA |
|---|---|---|---|---|---|---|
| **NEURONAL CULTURES** | Parrini et al., Molecular Terapy,2021 [1] | Hippocampal neurons | 16–20 DIVs | GABA (100 µM) | Cell-attached patch clamp | no |
| **BRAIN SLICES** | Stein et al., Journal of comparative neurology,2004[2] | hippocampal pyramidal neurons | P 14 | // | Gramicidin-perforated patch clamp | Only average data |
| | He et al., Journal of Neuroscience, 2014[3] | layer IV neuron in the somatosensory cortex | P 15 | // | | Only average data |
| | Wang et al., Brain Structure and Function, 2017[4] | layer II cortical neurons | 6-week-old | // | | Only average data |
| | Tyzio et al., Epilepsia, 2007 [5] | CA1 pyramidal neurons | P 25 | Isoguvacine (10 µM) | | no |
| | Tyzio et al., Science, 2014[6] | CA1 pyramidal neurons | P 30 | Isoguvacine (10 µM) | $GABA_A$R single channel currents | Only average data |
| | Yin et al., Toxicology Letters, 2016 [7] | VCN neurons | P 15-19 | BIC (50 µM) | Cell attached patch clamp | Only average data |
| | Dargaei et al., PNAS, 2018 [8] | CA1 pyramidal neurons | 4-5 weeks | GABA (100 µM)/ BIC (20 µM) | | no |
| | Lozovaya et al., Science report, 2019[9] | CA3 pyramidal neurons | P 15 | Isoguvacine (10 µM) | | no |
| | Parrini et al., Molecular Terapy, 2021[10] | CA1 pyramidal neurons | 8 -12-week-old | BIC (20 µM) | | no |
| | Khazipov et al.,2004[11] | CA1 pyramidal neurons | P 35 | Isoguvacine (10 µM) | Extracellular field potential | no |
| | Tyzio et al., Epilepsia, 2007 [5] | CA1 pyramidal neurons | P 25 | Isoguvacine (10 µM) | | no |

| | | | | | | |
|---|---|---|---|---|---|---|
| | Lysenko et al., Neurobiology of Disease,2018[12] | CA3 pyramidal neurons | P 22 | Isoguvacine (10 µM) | | no |
| | Pisella et al., Science Signaling, 2019[13] | CA3 pyramidal neurons | P 15-20 | Isoguvacine (10 µM) | | no |
| ***IN VIVO*** | Alfonsa et al.,2023 | L2/3 pyramidal neurons | 8 weeks old | Light activation of ChR2-expressing interneurons | Gramicidin perforated recordings | no |
| **iPSC-derived NEURONS** | Tang et al., PNAS, 2016 [14] | iPSC NEURONS | 2-3 months | GABA (100 µM) | Gramicidin-perforated patch clamp | Only average data |

**Supplementary Table 2. Articles in the literature that did not mention any population of mature neurons with depolarizing responses to $GABA_AR$ agonists or only showed average data.**

Abbreviations: DIV, Days *in vitro*; P, postnatal day; VCN ventral cochlear nucleus; ChR2 channelrhodopsin-2.
